# Single-cell quantification of senescence burden reveals cell type-specific ageing dynamics across organs

**DOI:** 10.1101/2025.11.14.688272

**Authors:** U. Cherqui, I. Sopher, H. Akiva, O. Menahem, E. Kopitman, R. Blecher-Gonen, H. Keren-Shaul, N. Rachmian, A. Mayo, U. Alon, H. Gal, V Krizhanovsky

## Abstract

Cellular senescence, a hallmark of ageing, drives tissue dysfunction by promoting inflammation and fuelling disease. Yet, the dynamics of senescent cell accumulation across tissues and their cell type identity remain poorly understood. Here, we introduce the first, single-cell, protein-level approach, combining multiple senescence markers for the identification and quantification of senescent cells across multiple tissues in mice and in human PBMCs. Applying this method, we reveal widespread but heterogeneous changes in senescence marker expression across cell types and tissues. The cells we identify as senescent displayed transcriptomic senescence signatures, providing a direct molecular link between protein- and mRNA-level detection of senescence. Importantly, senescence accumulation was strongly coordinated within organs but showed little correlation across them, supporting the idea of a tissue specific progression of ageing. These findings refine our understanding of the tissue-specific dynamics of senescence accumulation with age, and provide a framework for evaluating diverse therapeutic interventions.

## Introduction

Ageing is a primary risk factor for a wide range of age-related diseases, including certain cancers, cardiovascular dysfunction and neurodegenerative disorders^1^. Thus, understanding the mechanisms regulating ageing has become a major interest in medical research. Amongst the many hallmarks of ageing, senescent cells (SnCs) accumulation with age is a key driver of tissue degeneration and age-related pathology^1,2^. Cellular senescence is a stable cell cycle arrest mechanism which is triggered by stress or damage and serves as a protective mechanism to limit the proliferation of potentially damaged cells^2,3^. This state is characterized by a failure to re-enter the cell cycle upon mitogenic stimuli, resistance to cell death, and persistent DNA damage response, driving the secretion of factors, such as chemokines, cytokines, microRNAs, growth factors and proteases, in a phenomenon called the senescence-associated secretory phenotype (SASP)^4^. Cellular senescence serves an essential role in physiological processes, including limb patterning, placenta function and tissue remodelling during embryonic development^5–7^, preventing excessive cell proliferation following damage or oncogene activation^8–10^ and promoting tissue repair^11–13^. However, in the long-term, the accumulation of SnCs ultimately causes chronic inflammation and slows regeneration, at such levels that it results in tissue ageing, tumorigenesis and various age-related disorders^2^.

Despite growing recognition of their pathological roles, the tissue- and cell-type-specific dynamics of SnCs accumulation *in vivo* remain poorly understood. It is unclear whether all organs and cell types accumulate SnCs uniformly with age, or whether this process exhibits tissue- or cell-type-specific patterns. A major barrier to resolving this question has been the lack of robust methods for identifying and quantifying SnCs at the single-cell resolution *in vivo*^14,15^. Indeed, SnCs exhibit a complex phenotype but lack a universal and specific marker. The most widely used method to identify SnCs for the last 30 years has been the SA-β-Gal staining technique^16^. However, an important limitation of this method is that it offers limited specificity and quantification possibilities. Transcriptomic approaches, including single-cell RNA sequencing (scRNAseq), face challenges due to the low abundance of SnCs and the inconsistencies between the mRNA and protein expression of senescence markers. Additionally, the use of single markers of cell cycle arrest, such as CDK inhibitors p16 (CDKN2A) or p21 (CDKN1A), in flow cytometry, has limitations, as these markers might be present in non-SnCs^14,15,17^.

To address the challenges of SnCs identification, guidelines for detecting SnCs *in vivo* with high confidence were recently published^14, 15^. In concordance with these guidelines, we developed the first, single-cell, protein-level flow cytometry approach, combining multiple senescence associated markers for the identification and quantification of SnCs across multiple tissues in mice. A three-dimensional analysis using this approach demonstrated the necessity of multi-marker profiling to identify SnCs. By recovering and profiling the lungs, liver, intestine, and blood from young and aged mice, we show that most cell types acquire a differential expression of senescence markers. Additionally, our data reveals that the abundance of SnCs varies markedly across tissues and cell types with age. Subsequently, scRNAseq on sorted immune and non-immune lung cells, triple-positive for the three selected senescence markers, shows that these cells also harbour transcriptomic hallmarks of senescence. Together, these results provide the first direct molecular link between protein-level detection and transcriptional signatures of senescence. Finally, correlation analysis reveals that while senescence burden is poorly correlated across organs, strong intra-organ correlations exist among distinct cell types. Overall, these findings reveal substantial heterogeneity in senescence burden across cell types and organs. They also identify the cell types that contribute to the accumulation of SnCs with age and provide a framework for a quantitative and cell-type specific approach for assessing senescence burden in whole organisms.

## Results

### Mice exhibit a cell-type dependent differential expression of senescence markers with age

The accumulation of SnCs has important implications towards an individual’s healthspan^2,18,19^. However, the amount of SnCs in different organs and cell types remains unknown, due to the challenges to identify and quantify SnCs in tissues. This issue has led researchers in the field to summarize targeted recommendations on how to identify these cells^14,15^. The overreaching conclusion is that a combination of markers is necessary to identify SnCs *in vivo*. To attempt the identification and quantification of SnCs in tissues in line with these recommendations, we selected three senescence markers, p16, γH.2ax and Bcl-xl^2,14,15^. These markers are each associated with a different phenotypic characteristic of SnCs: cell cycle arrest (p16), persistent DNA damage response (γH.2ax) and resistance to apoptosis (Bcl-xl)^2^. We used 15 young (3-4 months old) and 15 old (23-24 months old) female mice from which we recovered the lungs, liver, intestine and blood. We obtained a single-cell suspension of these organs and stained them with lineage markers for cell types identification, as well as the three selected senescence markers (γH.2ax, p16 and Bcl-xl, (3SMs)) for Flow cytometry analysis (Fig. 1a). We first studied the changes in the overall abundance of different cell types with age. We measured the proportion of resident and immune cell types in each organ and age group, to observe the changes in abundance of each cell population with time (Extended Data Fig. 1a). Among these changes, we observed a decrease in the fraction of NK cells and Alveolar Macrophages (AM) in lung tissues from aged mice (Extended Data Fig. 1a, b). We also observed a decrease in the amount of CD4 T cells in the blood, an increase in the percentage of CD8 T cells in the liver, as well as an increase in macrophages in the intestine of old mice (Extended Data Fig. 1a, b). The comparison of these results with a study^20^ which used cytometry-based high parametric analysis of the changes in immune populations in aged mice across various tissues, revealed marked similarities between the observations of both studies. Therefore, this experimental set up leads to results consistent with the literature, and we can use it for identification and quantification of SnCs in these specific cell types.

**Figure 1:**
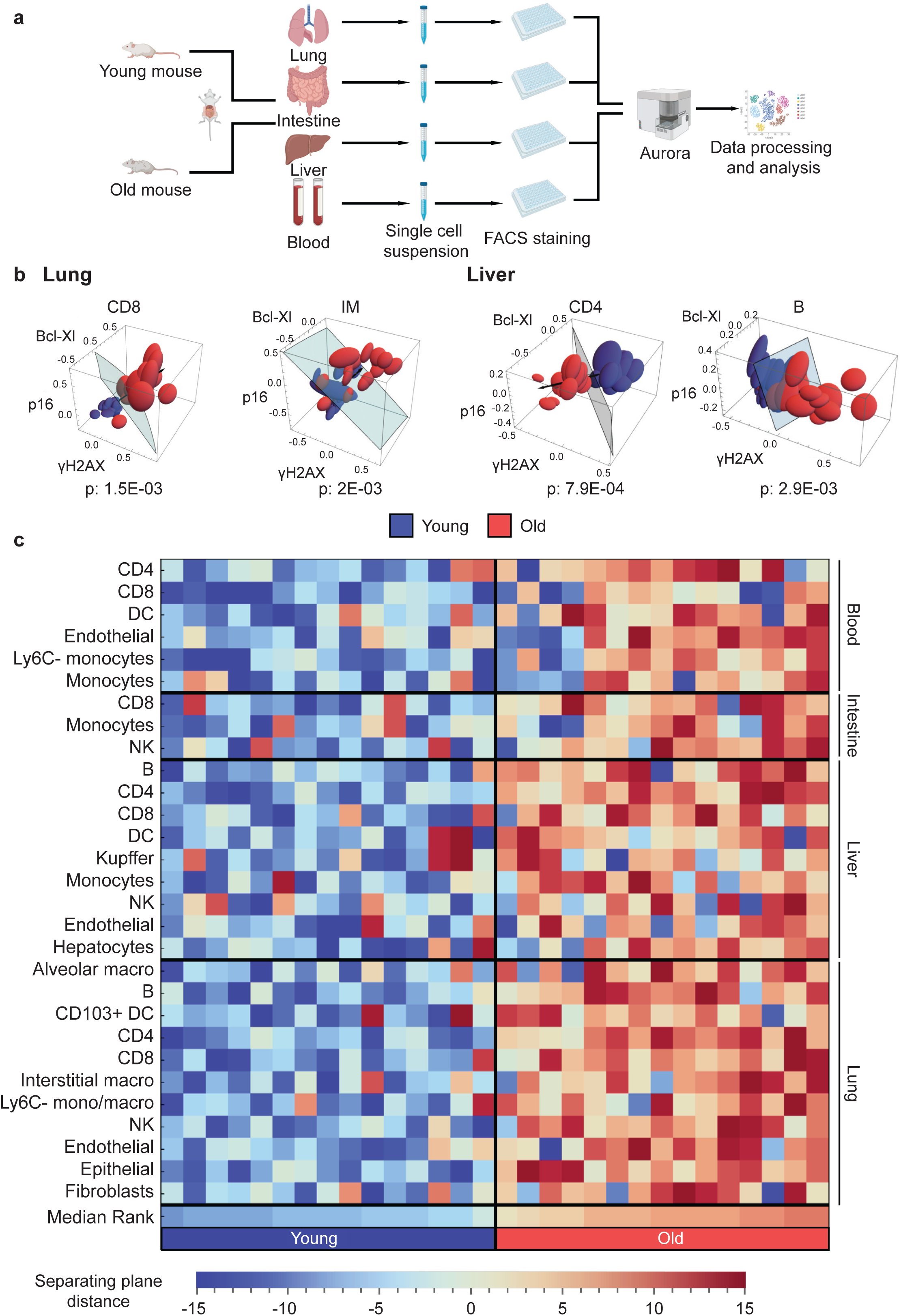
Combination of selected senescence markers distinguishes between young and old mice in multiple organs and cell types. **(a)** Schematic representation of tissues collection and staining for senescence quantification. Lung, liver, intestine and blood are collected from young (3-4 months) and old (23-24 month) female mice. A single-cell suspension is obtained for each lung, liver and intestine. The samples are stained with lineage markers antibodies for cell type identification, as well as with the senescence markers: p16, γH.2ax and Bcl-xl and analysed by flow cytometry. **(b)** A three-dimensional analysis using the 3SMs, helps define a ‘new’ composite senescence score based on the expression of these markers, and define a plane which can separate young from old mice in each cell type. This analysis automatically finds the best separating manifold by using a support vector machine (SVM) to define a separating plane and a separating direction between young and old groups. Lung CD8 T cells and interstitial macrophages (IM), as well as liver CD4 T and B cells are used as representative examples. **(c)** Heatmap of mice ranks for cell types which were significantly separated by the SVM, as exemplified in (b). Results from the three-dimensional analysis are projected on the vector directing young to old groups and the distance of each mice from the separating plane is calculated for each cell type. The blue colour indicates that the mouse fell on the ‘young’ part of the plane, while the red colour indicates that the mouse fell on the ‘old’ part of the plane. The intensity of the colour indicates the distance of the mouse from the separating plane. The division between the chronological age groups the mice belong to is indicated at the bottom.

Using this setup, we acquired data for protein expression of the 3SMs for all the organs, allowing us to study the expression of these markers in all cell types in these organs. To understand if a differential expression of the 3SMs can distinguish between young and old mice in different cell types in an unbiased manner, we set to test if we could use them to separate mice between age groups based on their expression levels alone. We used a three-dimensional analysis of these 3SMs to define a composite senescence score based on the ungated data of expression of these markers, for each cell type for each mouse, and defined a plane which could separate young from old mice in each cell type (Fig. 1b and Extended Data Fig. 2). This three-dimensional analysis allows us to find automatically the best separating manifold by using a support vector machine (SVM) to define a separating plane and a separating direction between young and old groups. This scoring system allows us to find in an unbiased manner, the cell types in which a significant difference exists between young and old groups based on the expression of the 3SMs. For example, CD8 T cells and interstitial macrophages (IM) in the lung, as well as CD4 T and B cells in the liver, have a clear separation between young and old mice (Fig. 1b). Out of the 43 cell types we obtained across organs, this approach could separate young from old mice in 29 cell types, in a statistically significant manner (Fig. 1b, c and Supplementary Tables 1 and 2), showing that many cell types across tissues develop changes in the expression of the 3SMs with age.

To obtain a more comprehensive view of the cell types which showed a significant, differential expression of the 3SMs with age, we used the results from this three-dimensional analysis to project each mouse from each cell type on the vector directing from young to old groups and made a heatmap of mice ranks per cell types (Fig. 1c). We observed that for these cell types, most young mice fall in the ‘young’ part of the plane, while most old mice fall in the ‘old’ part of the plane, showing a clear distinction between the two age groups (Fig. 1c). Together, these results show that most, but not all cell types acquire a differential expression of senescence markers with age.

### Multiple markers are necessary to identify senescent cells

The use of multiple markers is necessary for valid SnCs identification^14, 15^, and it indeed allowed us to distinguish young from old mice in numerous cell types and tissues in an unbiased manner. However, the relative contribution of single markers and their combination was not estimated before quantitatively. Therefore, we aimed to study the relative contribution of multiple markers to accurately study the dynamics of SnCs during ageing. When calculating the fraction of cell types which can be separated between young and old mice using the 3SMs, the use of multiple markers allows to separate more cell types accurately, with two markers being better than one and three markers being better than two (Fig. 2a). Additionally, we looked at the contribution of each marker in defining the plane that separates young from old mice in each cell type and found that at least two markers play a role in every cell type, with three markers being necessary in most cases (Fig. 2b). We then selected cell types with different levels of contribution from the 3SMs in defining this plane, to verify whether the 3SMs still played an important role in explaining the variance in those cell types. Intestinal macrophages showed low p16 contribution, while lung B cells showed important contribution from all 3 markers and liver neutrophils showed low γH.2ax contribution (Fig. 2b). We calculated their principal component analysis (PCA) loadings to determine the impact of each senescence marker on the variance observed to separate young from old mice. We found that even when a marker only showed limited contribution in defining the plane, it played an important role in the variance between mice (Fig. 2c). Together, these results experimentally confirm the necessity of using multiple markers to study cellular senescence and reveal that relative contribution of every marker is cell type specific.

**Figure 2:**
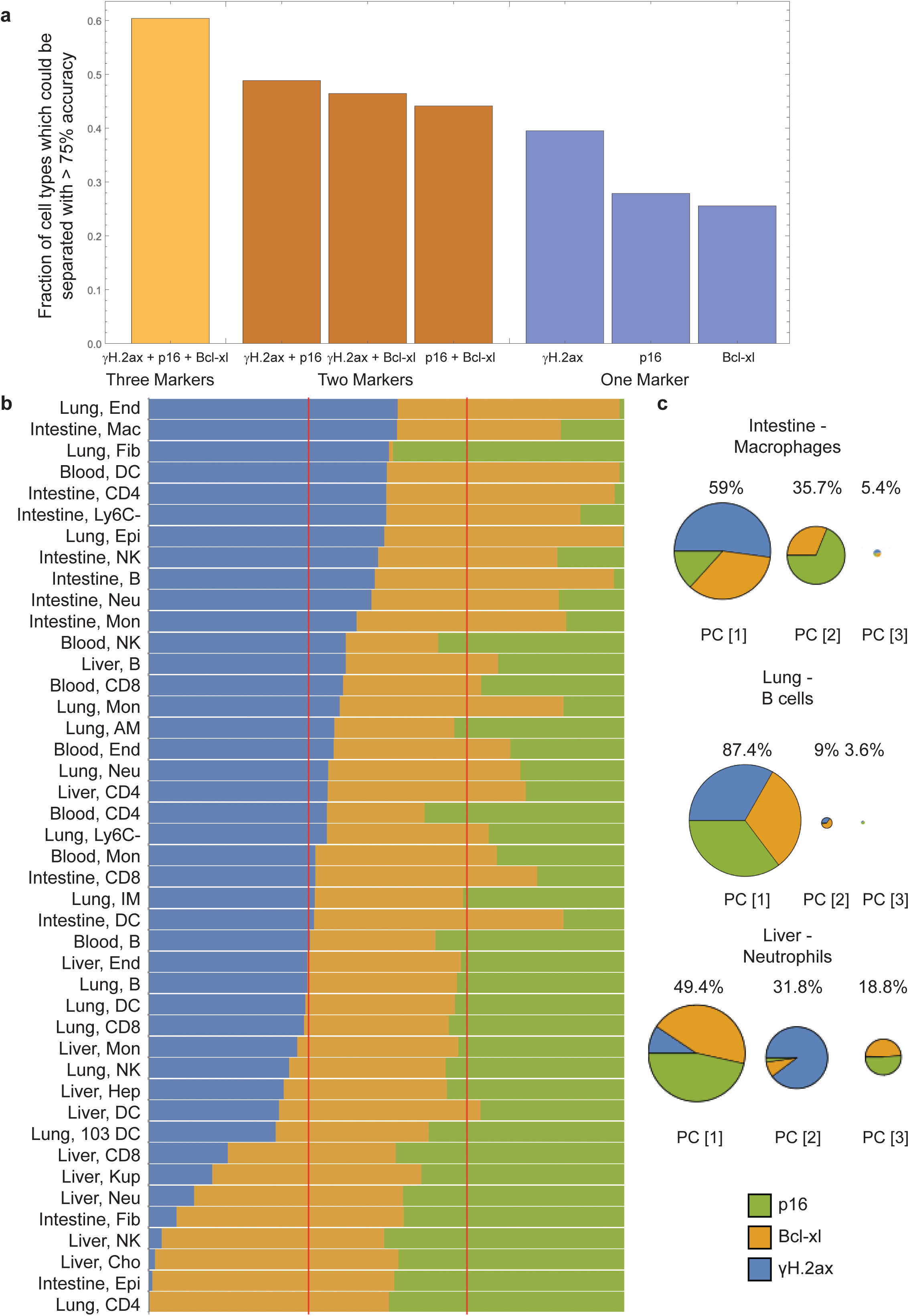
Multiple markers are necessary to study senescent cells in tissues. **(a)** The fractions of cell types which can be separated between young and old mice with over 75% accuracy based on the three indicated senescence markers or different combinations of two senescence markers, or each marker alone. **(b)** The contribution of each marker in defining the plane that separates young from old mice in each cell type (Fig. 1b). The cell types and organs are indicated on the left. Epi- Epithelial cells; End- endothelial cells; Fib- fibroblasts; Hep- hepatocytes; Cho- cholangiocytes; B- B cells; CD4- CD4 T cells; CD8- CD8 T cells; NK- NK cells; AM- alveolar macrophages; Kup- Kupffer cells; 103 DC- CD103+ DCs; Neu-Neutrophils; IM- Interstitial macrophages; DC- dendritic cells; Mon- monocytes; Ly6C- -Ly6C-monocytes; Mac- macrophages. **(c)** Cell types with different levels of contribution from the 3SMs in defining the plane separating young from old mice were then selected based on results from (b). Intestinal macrophages showed low p16 contribution, while lung B cells showed important contribution from all 3 markers and liver neutrophils showed low γH.2ax contribution. The PCA loadings calculating the impact of each senescence marker on the variance observed was then calculated for these cell types, demonstrating that in all cases, all three markers are needed to explain most of the variance. PC[1]: Principal component 1; PC[2]: Principal component 2; PC[3]: Principal component 3.

### Senescent cell burden differs between organs and cell types

The combination of 3SMs separates samples from young and old mice in most cell types. To study which cell types and tissues accumulate SnCs with age, we used these markers to identify and quantify SnCs. In order to quantify SnCs, we used a manually curated gating strategy to calculate the proportion of cells expressing all 3SMs, which we named SenM+ cells (senescence markers positive cells), in each cell type of the lung, liver, intestine and blood (Fig. 3a-d).

**Figure 3:**
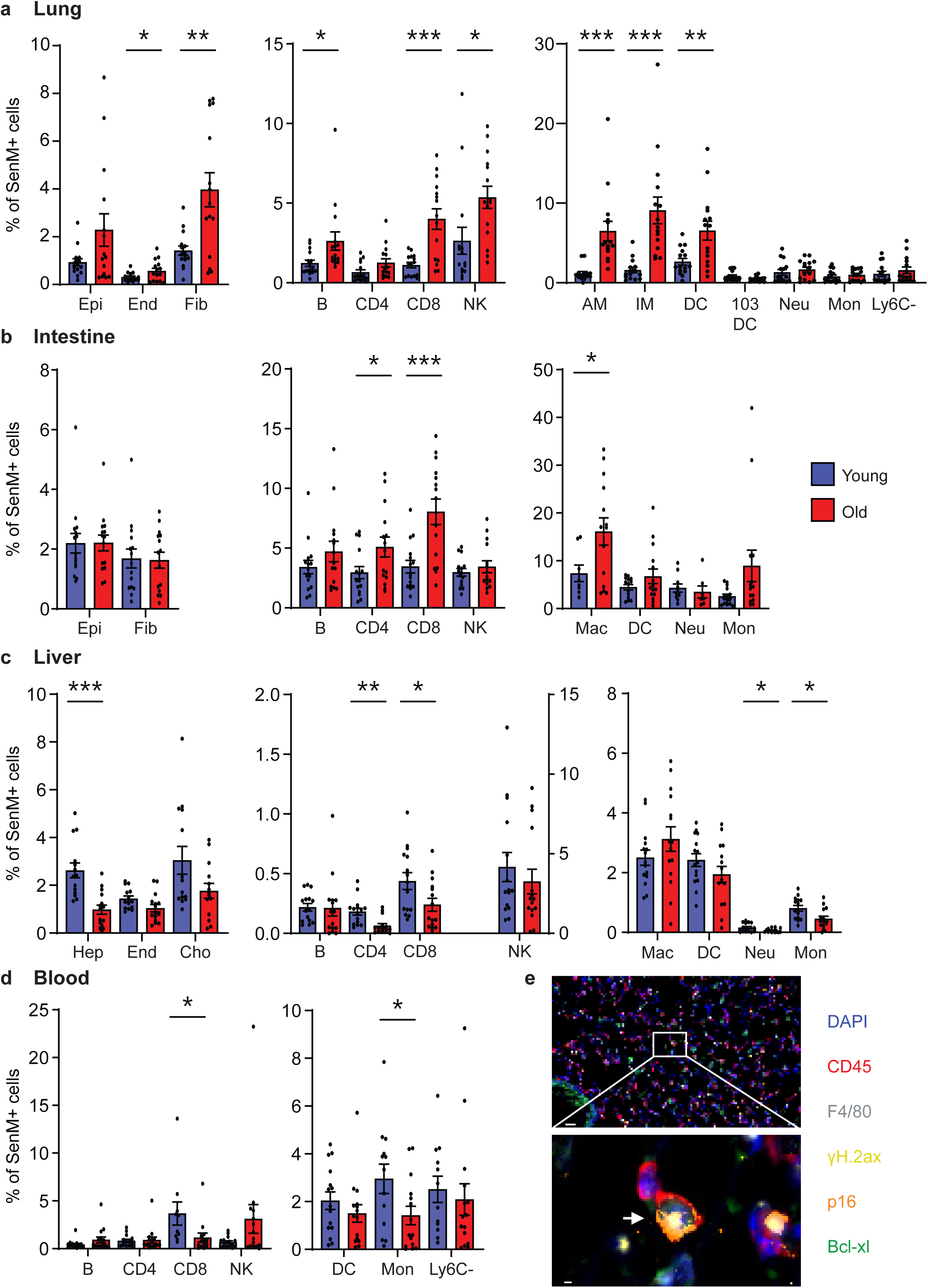
The percentage of cells positive for the selected senescence markers (SenM+) for each cell type of young and old mice. The quantification is presented for each organ separately as follows: **(a)** lung **(b)** liver **(c)** intestine **(d)** blood. Quantification of the SenM+ populations was obtained via gating on Spectroflow (see Extended Data Fig. 1, 4). Only triple positive cells for p16, γH.2AX and Bcl-xl were considered SenM+. A P value < 0.05 was considered statistically significant, following an unpaired t-test analysis (* < 0.05, ** < 0.01, *** < 0.005), error bars indicate SEM. Resident, lymphoid and myeloid cells are shown on different panels. Epi- Epithelial cells; End- endothelial cells; Fib- fibroblasts; Hep-hepatocytes; Cho- cholangiocytes; B- B cells; CD4- CD4 T cells; CD8- CD8 T cells; NK- NK cells; AM-alveolar macrophages; Kup- Kupffer cells; 103 DC- CD103+ DCs; Neu- Neutrophils; IM- Interstitial macrophages; DC- dendritic cells; Mon- monocytes; Ly6C- -Ly6C- monocytes; Mac- macrophages. **(e)** Multispectral imaging of lung tissues from young and old mice using the PhenoImager HT system with antibodies directed to CD45 and F4/80 to identify macrophages and antibodies directed to our selected senescence markers, γH.2ax, p16 and Bcl-xl. The arrow points towards a cell positive for every marker. Scale bars, 20µm and 2µm.

In lung tissues, we observed an increase in the percentage of SenM+ cells in old mice in various resident and immune cell types (Fig. 3a). We observed a significant 2.8-fold increase in fibroblasts (Fib, p=0.0018), as well as a 1.8-fold increase in endothelial cells (End, p=0.0454) (Fig. 3a). In the immune system, the components of both innate and adaptive arms accumulate SenM+ cells. We observed a significant 3.6-fold increase in CD8 T cells (p=0.0002), a 2.1-fold increase in B cells (p=0.0291) and a 2-fold increase in NK cells (p=0.0191). Significant increases were also observed in macrophages and dendritic cells, with a 5.5-fold and a 5.7-fold increase in alveolar (AM, p=0.0002) and interstitial (IM, p=0.0002) macrophages respectively, as well as a 2.4-fold increase in dendritic cells (DC, p=0.0044) (Fig. 3a). Therefore, multiple cell types in the lung accumulate SnCs with age, which could impact the organ’s function.

We observed a significant increase in the percentage of SenM+ macrophages in lung tissues in old mice. We therefore aimed to identify these cells in tissues using immuno-histochemistry. We used the Opal Multiplex IHC Assay and stained lung tissues from young and old mice with antibodies directed to CD45 and F4/80 to identify macrophages and antibodies directed to the 3SMs, γH.2ax, p16 and Bcl-xl. We successfully detected senescent macrophages expressing all 3SMs (Fig. 3e and Extended Data Fig. 3). Therefore, 3SM-positive cells can be identified in tissues *in situ*.

In the intestine, we have not identified an increase in SenM+ cells in the resident cell populations we could recover, while we did detect it in some of the components of its immune system. T cells and macrophages show a significant increase in the proportion of SenM+ cells with age. Indeed, a 1.7-fold and a 2.3-fold increase in the percentage of SenM+ cells is observed in CD4 (p=0.0367) and CD8 (p=0.0006) T cells respectively, while a 2.2-fold increase can be observed in macrophages (Mac, p=0.0424) (Fig. 3b). Overall, while some of the immune cells in the intestine accumulate senescent cells with age, the trend is not as widespread as in the lung.

We then analysed the accumulation of senescent cells with age in the liver. Surprisingly, in contrast to the lung and intestine, no increase in senescence burden could be observed in any of the cell types we recovered from in the liver of old mice. Of note, a significant decrease in the amount of SenM+ cells could be observed in hepatocytes (Hep, 2.7-fold; p=0.0001), as well as CD4 (2.9-fold; p=0.0015) and CD8 (1.8-fold; p=0.0329) T cells, neutrophils (Neu, 2.5-fold; p=0.0273) and monocytes (Mon, 1.8-fold; p=0.0105) (Fig. 3c). A limitation for the hepatocytes analysis stems from the fact that only diploid hepatocytes could be analysed using this method, while the proportion of multiploid hepatocytes is known to increase with age^21^. The absence of increase in senescence burden in liver cells in old mice may be explained by the liver high regenerative ability and immunotolerance^22,23^. Finally, in the blood, no cell type of hematopoietic origin showed an increase in senescence burden with age (Fig. 3d). However, a significant 8-fold decrease in SenM+ cells can be observed in circulating endothelial cells (CECs, p=0.0020), as well as a significant decrease in CD8 T cells (3.2-fold; p=0.041) and monocytes (Mon) (2.1-fold; p=0.0394) (Fig. 3d and Extended Data Fig. 4a). The half-life of immune cells in the blood is much shorter than in organs, which may explain why no significant increase in SnCs could be observed. To achieve a comprehensive view of the cell type and organ specific changes in senescence burden with age, we made a heatmap of the ratio of the percentage of SenM+ cells in old over young mice for each cell type (Extended Data Fig. 4b). This heatmap underscores that the lungs and intestine of mice seem to develop a high senescence burden with age, while very few changes can be observed in the liver and the blood.

One of the limitations from this analysis stems from the bias associated with the manual gating of the SenM+ populations. To further validate our results, we selected the cell types from the unbiased three-dimensional analysis based on the ungated data, in which the vector defining the separating plane between young and old mice moved in a direction where all of the 3SMs went up in aged mice (Extended Data Fig. 4c). We then calculated their percentage of SenM+ cells in an unbiased manner, by applying a threshold on the expression distribution of the 3SMs, above which cells were considered SenM+ (Extended Data Fig. 5d). Using this method, lung and intestine immune cell types showed a significant increase in senescence burden with age (Extended Data Fig. 4d), while no liver or blood cell type showed such an increase. These results support the notion that an increase in senescence burden occurs in lung and intestine tissues of aged mice, unlike in liver or blood. Overall, these results indicate that different cell types and organs develop distinctive senescence burden with age.

### SenM+ cells exhibit a senescent transcriptional signature

We developed a method to identify SnCs at the single-cell level using protein-based markers in flow cytometry. To study the nature of the cells we identified by their comprehensive mRNA expression profile and compare it to published profiles and signatures, we aimed to collect these cells and analyse them using single-cell RNA sequencing (scRNAseq). We collected cells from lungs of 24-months old mice and stained these cells with CD45 to identify immune and non-immune cells, along with the 3SMs - γH.2ax, p16 and Bcl-xl. We then sorted cells co-expressing these markers into senescent immune (SenM+;CD45+) and non-immune (SenM+;CD45-) cells. Finally, we used the Chromium Fixed RNA-profiling for multiplexed samples (10X Genomics) approach to perform scRNAseq on sorted cells. We then compared this dataset with ‘total lung’ scRNAseq data obtained from Gene Expression Omnibus (GEO, deposited as GSE141259)^24^ (Fig. 4a-d). Using canonical cell type markers, we identified multiple non-immune (Fig. 4a and Extended Data Fig. 5a) and immune (Fig. 4c and Extended Data Fig. 5c) cell types to compare to total lung data. A major challenge in senescence transcriptomics lies in the phenotypic heterogeneity and context-specific signatures of SnCs in different cell types and tissues. To address this, we used the open-source Python package SenePy which was recently developed^25^ to compute senescence scores across the identified cell types. We observed markedly elevated senescence across SenM+;CD45-(Fig. 4b) and SenM+;CD45+ (Fig. 4d) populations. Analysis of the SenePy score revealed a significant increase in the proportion of SnCs in every SenM+;CD45- (p<0.0001, Fig. 4e and Extended Data Fig. 5b) and SenM+;CD45+ (p<0.02, Fig. 4f and Extended Data Figure 5d-f) cell type analysed, relative to control. These results further confirm that cells identified by co-expression of γH2AX, p16, and Bcl-xl at the protein level, harbour transcriptomic hallmarks of senescence. Together, these results provide a direct molecular link between protein-level detection and transcriptional signatures of senescence, demonstrating that cells expressing high levels of γH.2ax, p16 and Bcl-xl, exhibit a senescent transcriptomic signature at single-cell resolution in both immune and non-immune cell types.

**Figure 4:**
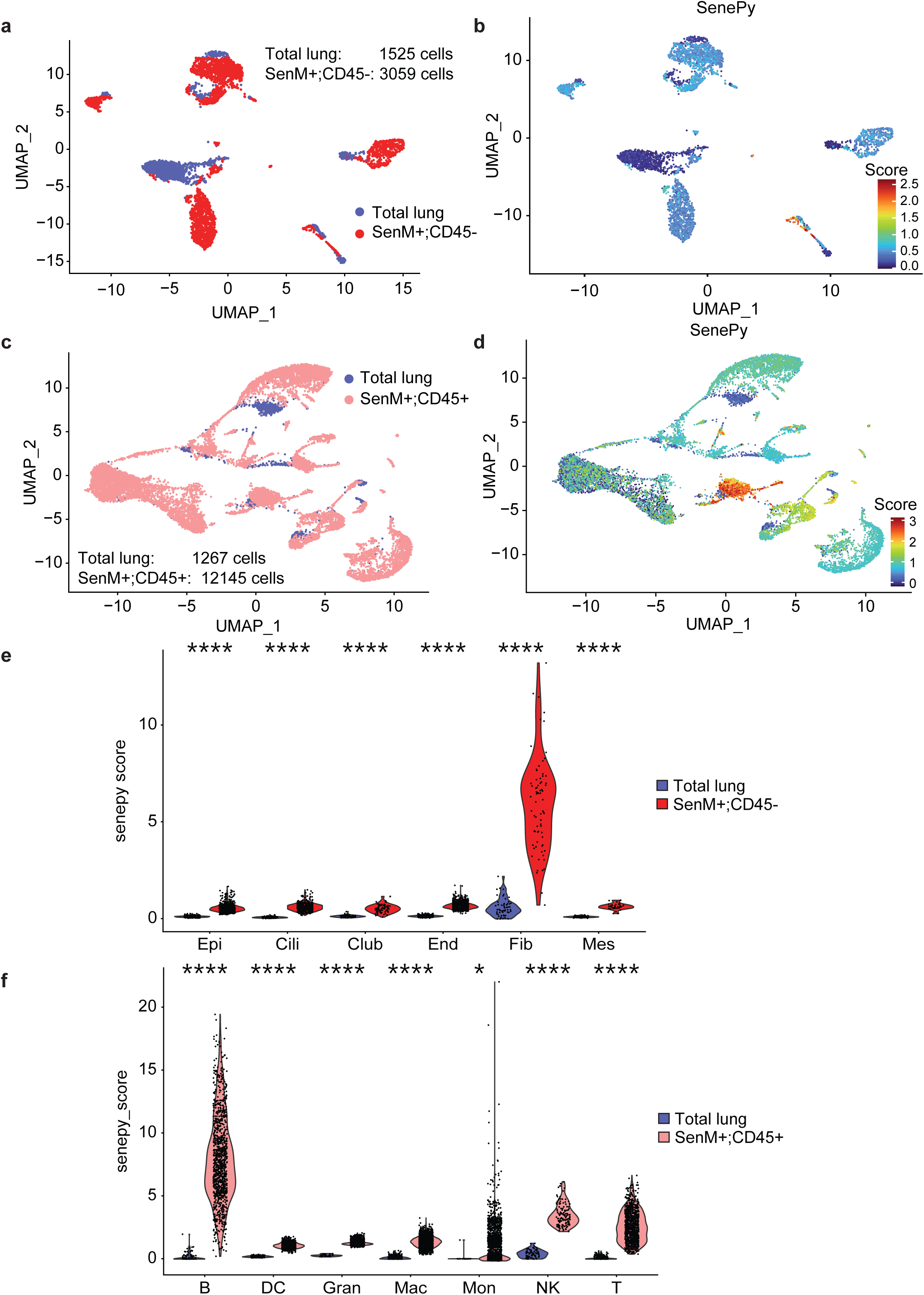
Analysis of scRNAseq of SenM+ cells. Lungs from 24-months old mice were collected and dissociated. These cells were then stained with CD45 to identify immune and non-immune cells, along with selected senescence markers - γH.2ax, p16 and Bcl-xl, to identify SenM+ cells. Then, SenM+;CD45+ and SenM+;CD45- were sorted for scRNAseq. The cells were sequenced using the Chromium Fixed RNA-profiling for multiplexed samples (10X Genomics) approach. SenePy score in SenM+ cells in different cell types was then compared to matched total lung cell types. **(a)** UMAP representing SenM+;CD45- and total lung CD45- cells. **(b)** SenM+;CD45- and total lung CD45- cells scored using the SenePy signature. The UMAPs color scale was created using a log1p(senepy_score). **(c)** UMAP representing SenM+;CD45+ and total lung CD45+ cells. **(d)** SenM+;CD45+ and total lung CD45+- cells scored using the SenePy signature. The UMAPs color scale was created using a log1p(senepy_score). **(e)** Violin plots of SenePy scores based on the universal SenePy signature in SenM+;CD45- and total lung CD45- cell types (The cell-type-specific SenePy signature was used for fibroblasts, as the only applicable cell type). **(f)** Violin plots of SenePy scores based on the cell-type-specific SenePy signature in SenM+;CD45+ and total lung CD45+ cell types. A P value < 0.05 was considered statistically significant, following a Wilcoxon test analysis (* < 0.05, ** < 0.01, *** < 0.001, **** < 0.0001).

### Senescence burden correlates within organs

Accumulation of SnCs in a tissue could represent a parameter of biological ageing, specific to the tissue. We then asked if this accumulation is correlated between different tissues in the organism. We first calculated the overall senescence burden of each organ in each mouse. The cumulative analysis of senescence burden in all tissues showed marked variability between individual mice but showed that cumulative burden is increased in the lungs and intestine of old animals (Fig. 5a and Extended Data Fig. 6). Additionally, this analysis underscored the variability in the presence of SnCs in different cell types and organs. To understand if different organs accumulate SnCs at the same rate, we performed a correlation analysis to ask whether mice with higher senescence in specific organs were more likely to exhibit higher senescence levels in other organs (Fig. 5b). The correlation analysis comparing the overall senescence burden between each organ showed that a significant inverse correlation between the lung and liver (p=0.025), as well as blood and liver (p=0.043) existed (Fig. 5b). However, we observe that these were both weak correlations (lung/liver R^2^: 0.3306; blood/liver: R^2^: 0.2794). This proposes that different organs mostly accumulate SnCs independently of each other and that the amount of SnCs in specific organs or cell types may better define ageing of a specific organ than their general accumulation in the body.

**Figure 5:**
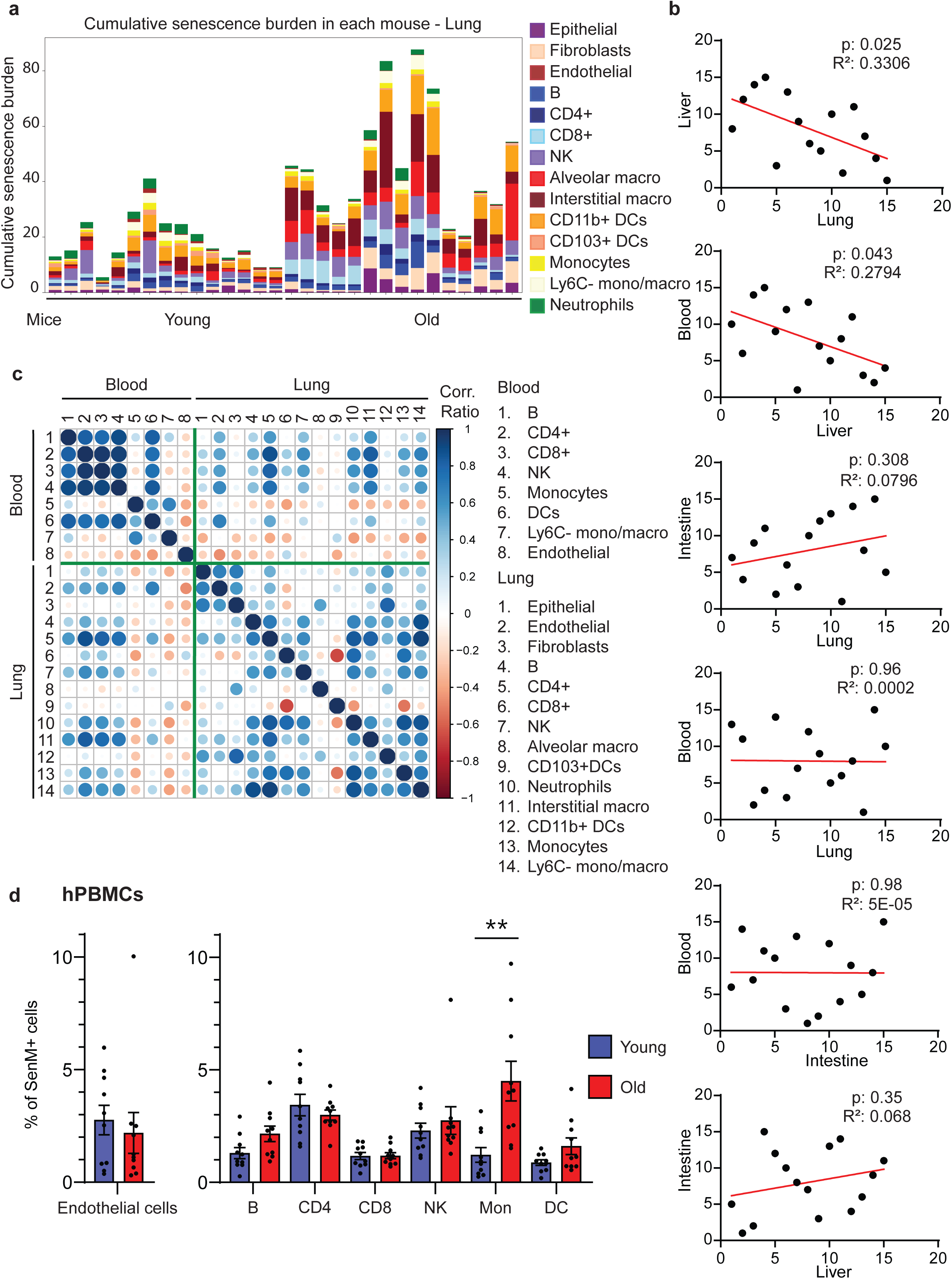
SenM+ cell burden in different organs and cell types in mice and in human PBMCs. **(a)** The cumulative senescence burden of each mouse was calculated by adding together the percentages of senescent cells in each of their cell types. Each bar represents a single mouse. **(b-c)** Correlation analysis of SenM+ cell burden between different organs and cell types was performed in young and old mice **(b)** Mice were ranked based on their overall senescence burden for each organ. We subsequently calculated the correlation between their ranks across organs. The organs correlated are indicated at the axes. The R^2^ score was calculated via simple linear regression using GraphPad Prism. **(c)** A correlation plot comparing each cell types in the lung and blood of aged mice, was created using R and RStudio with “corrplot” libraries. **(d)** Quantification of the SenM+ levels in hPBMCs from young and old donors was obtained via manual gating on Spectroflow (see Extended Data Fig. 11). Triple positive cells for p16, γH.2AX and Bcl-xl were considered as SenM+. The following cell types were evaluated: Endothelial cells; B- B cells; CD4-CD4 T cells; CD8- CD8 T cells; NK- NK cells; Mon- monocytes; DC- dendritic cells. A P value < 0.05 was considered statistically significant, following an unpaired t-test analysis (* < 0.05, ** < 0.01, *** < 0.005), error bars indicate SEM.

To compare the senescence burden of different cell types within and across organs we created correlation plots between them (Fig. 5c and Extended Data Fig. 7 and 8). These results show that a correlation between different cell types exists within organs, demonstrating that while organs may accumulate SnCs independently of each other, different cell types within these organs seem to accumulate senescence concurrently (Fig. 5c and Extended Data Fig. 7 and 8). On the other hand, cell types across organs do not seem to accumulate SnCs synchronously, as little correlation could be observed, even when comparing similar cell types (Fig. 5c and Extended Data Fig. 7 and 8). Therefore, different cell types accumulate SnCs concomitantly within organs, while similar cell types do not seem to accumulate SnCs concurrently across organs.

### Quantification of senescence levels in human PBMCs

Following the study in mice, we aimed to adapt this method to quantify SenM+ cells at the single-cell level using molecular markers on human samples. We obtained human PBMCs from ten young (<35 years old) and ten old (>65 years old) healthy male donors. We stained these cells with lineage markers, as well as the 3SMs (γH.2ax, p16 and Bcl-xl) and analysed these cells via flow cytometry. Owing to the gating strategy used for the mouse analysis which identified SenM+ cells as senescent by their expression profile, we used a similar gating strategy to identify SenM+ cells in human PBMCs and quantify the proportion of SenM+ cells present in each cell type. In these samples, we were able to quantify SenM+ cells in B-cells, CD4 and CD8 T-cells, NK cells, Monocytes, Dendritic cells and CECs. We found that in human PBMCs, from the above analysed cell types, only monocytes showed a statistically significant (p=0.0026) increase in the proportion of SenM+ cells, with a 3.7-fold increase with age (Fig. 5d). Of note, there is stronger variability in humans compared to mice, in both the abundance of different cell types between individuals and in the amount of SenM+ cells. Both types of variability could contribute to lack of significant differences between cells from young and old donors. Overall, these experiments demonstrate that SnCs burden can be quantified within specific PBMC populations of individual patients, paving the way for population-wide monitoring of cellular senescence.

## Discussion

A major barrier to understanding whether all organs and cell types accumulate SnCs homogenously with age, has been the lack of robust methods for identifying and quantifying SnCs at the single-cell resolution *in vivo* ^14, 15^. Here, we introduce the first, single-cell, protein-level flow cytometry approach which combines multiple senescence-associated markers to identify and quantify SnCs across multiple tissues. Applying this method in mice lungs, intestines, liver and blood, we show that most, but not all cell types in these organs acquire a differential expression of the senescence markers, p16, γH.2ax and Bcl-xl with age. Indeed, a three-dimensional analysis using these markers, distinguished young from old mice in most but not all cell types, underscoring the heterogeneity of senescence accumulation and the importance of single-cell resolution.

Our results reinforce recently established guidelines that emphasize the necessity of using multiple markers to detect senescence *in vivo*^14, 15^. This can be explained by the fact that no specific marker for SnCs has been identified. Indeed, the most used markers of senescence, such as p16, p21 or γH.2ax, have all been shown to be expressed in non-SnCs in other contexts^26–28^. We found that marker combinations increased discriminatory power between young and old mice, with three markers outperforming two, and two outperforming one. Although contributions from each marker varied by cell type, at least two markers were required in all cases, and three were necessary in most. All three markers were also necessary to explain most of the variance in our three-dimensional analysis. These findings also demonstrate that relative contribution of every marker is cell type specific, suggesting that optimal marker combinations may differ across cell types and tissues. Indeed, our study was limited to the use of three senescence-associated markers, which were each selected to reflect a different phenotype of SnCs: cell cycle arrest (p16), persistent DNA damage response (γH.2ax) and resistance to apoptosis (Bcl-xl). There is no certainty that these markers are the most adapted to study senescence in each of the cell types we analysed. Expanding beyond the three markers used here to include p21, p53, or SA-βGal may refine senescence profiling further and future work should focus on establishing the most effective marker combinations for distinct cell types and tissues.

By manually quantifying the proportion of triple-positive cells (SenM+), we aimed to identify the cells and tissues showing the greatest age-associated increases in senescence burden. We observed striking tissue specificity. Lung and intestinal cell types showed robust increases in senescence burden with age, whereas liver and blood cells did not. Ageing was associated with a significant rise in SenM+ fibroblasts and endothelial cells in the lung, a shift that could compromise tissue function and promote the development of different age-related diseases. Fibrotic lung diseases were for example shown to be mediated by senescent cells^29^. Several populations from both arms of the immune system exhibited increased proportions of SenM+ cells in the lung, most notably B, T and NK cells, as well as macrophages and dendritic cells. Most of these cells are believed to play important roles in clearing SnCs throughout life^13, 30, 31^, and their increased senescence burden in old age may underlie the age-associated decline in immune surveillance, and development of chronic inflammation. Similarly, in the intestine, T cells and macrophages both showed an increase in their proportion of SenM+ cells. Conversely, in the liver, hepatocytes exhibited a decrease in SenM+ cells with age, though interpretation is complicated by age-related ploidy shifts^21^. Indeed, multiploidy in hepatocytes increases with age, while only diploid cells could be studied. The lack of senescence accumulation in the liver was surprising but may reflect its regenerative and immunotolerant properties which make it less susceptible to many of the changes associated with ageing^22^. Moreover, studies have shown that p16-positive endothelial cells as well as hepatic stellate cells in the liver play an important role in limiting liver fibrosis^13, 23^. SnCs in the liver may therefore play an important role to ensure proper organ function. In the mouse blood, there was an absence of any age-associated increase in senescence burden, which may be explained by the short half-life of circulating immune cells. However, a major decrease in SenM+ cells percentage can be observed in CECs. Because endothelial cells normally safeguard vascular integrity yet detach after damage^32^, the observed decline in senescent CECs raises the possibility that these cells play an unrecognized role in maintaining vessel function or integrity. Together, these findings highlight the organ- and cell type-specific nature of senescence accumulation and argue for a more targeted approach in understanding its contribution to ageing and age-related disorders.

Our method provides a framework for testing senolytic interventions and for monitoring senescence dynamics following therapies. Tailoring senolytic treatments to specific cell types or organs may increase therapeutic benefit and mitigate cytotoxicity, thereby overcoming a major barrier to their clinical application. Expanding this method to diverse organs and multiple time points will not only identify the cell types and tissues most affected by senescence in old age but also elucidate the temporal dynamics of SnCs accumulation. One of the limitations from this experiment results from the bias associated with the manual gating of the SenM+ populations. To mitigate this, we used the three-dimensional analysis to identify cell types with age-associated increases in 3SMs and applied an unbiased thresholding approach to define SenM+ cells. This analysis confirmed our initial findings, with elevated senescence burden restricted to lung and intestinal cell types in old mice. We further validated the senescent identity of SenM+ cells. Immunohistochemistry confirmed the presence of lung macrophages expressing these three markers. Furthermore, scRNAseq on sorted SenM+ lung cells shows that these cells also harbour transcriptomic hallmarks of senescence across immune and non-immune lineages. Together, our results provide the first direct molecular link between protein-level detection and transcriptional signatures of senescence, highlighting that cells expressing p16, γH.2ax and Bcl-xl also display a senescent transcriptomic signature at single-cell resolution, in both immune and non-immune populations.

Finally, our findings inform the broader question of biological ageing. Chronological age does not capture the heterogeneity of health status, whereas biological age aims to do so. A central debate is whether a single, unified ageing clock can reliably capture overall biological age or whether organ-specific clocks are required to provide a more accurate representation of tissue-specific ageing. Using senescence burden as a marker for ageing, we observed that organs accumulate senescence largely independently of one another. This indicates that senescence burden within individual organs may more faithfully reflect ageing than aggregate measures across the body. Moreover, in our data, senescence burden was strongly correlated among diverse cell types within the same organ, yet largely uncorrelated across organs, even for analogous populations. Together, these findings indicate that the local tissue environment, rather than cell identity, is the dominant determinant of senescence dynamics.

In summary, this study introduces the first, single-cell, protein-level flow cytometry method, combining multiple senescence-associated markers for the identification and quantification of SnCs across multiple tissues in mice and in PBMCs in humans. Our findings reveal profound heterogeneity in senescence burden across cell types and organs and establish protein-level senescence markers as consistent with transcriptomic signatures. We also found coordinated senescence dynamics within organs but little correlation across them, highlighting the tissue specificity of senescence. Together, these findings advance the molecular toolkit for senescence and ageing biology, refine our understanding of its tissue-specific dynamics, and provide a foundation for testing seno-therapeutic strategies.

## Methods

### Mice

C57BL/6 female mice were obtained from Harlan Laboratories. Young mice were 3-4-months old, while old mice were 22-24-months old. The Weizmann Institute of Science Animal Care and Use Committee (IACUC, 03810525-2) approved all the animal studies described in this work.

### Organs dissociation to a single-cell suspension

Mice were anaesthetized with Pentobarbital Sodium (CTS Chemical Industries Ltd). Heparin was then injected through the mouse’s heart and blood was collected via cardiac puncture. Then, mice were perfused with cold PBS for blood clearance. The lungs, livers and intestines of mice were recovered, washed in PBS and minced. PBMCs were obtained from the blood by density gradient centrifugation, using Ficoll-Paque (Sigma-Aldrich Solutions). The lungs were incubated at 37°C on a shaker, in DMEM/F-12 supplemented with 1 mg/ml DNase I (Roche), 1 mg/ml collagenase type IV (Sigma-Aldrich Solutions) and 2.4 units/ml dispase II (Roche) for 25 minutes. The livers were incubated in similar conditions in DMEM/F-12 supplemented with 2mg/ml DNase I and 2mg/ml collagenase type IV for 20 minutes. The cell suspensions were then filtered through a 100-µm nylon mesh and centrifuged. The intestines were incubated in PBS supplemented with 0.01M EDTA (Sigma-Aldrich Solutions) and 0.0015M DTT (Sigma-Aldrich Solutions) for 20 minutes at 4°C. Subsequently, the tissues were moved into PBS supplemented with 2mg/ml DNase I and 0.01M EDTA and incubated on a shaker at 37°C for 10 minutes. The tissues were gently shaken and the cells recovered. The remains of the tissues were incubated in RPMI supplemented with 10% FBS, 100 units/ml of penicillin and 100 mg/ml of streptomycin, 0.03mg/ml DNase I, 0.2mg/ml collagenase type IV for 45 minutes at 37°C on a shaker. The cell suspensions were then filtered through a 100-µm nylon mesh and centrifuged. The red blood cells were lysed in each sample using Ack lysing buffer (Thermo Fisher Scientific). The cell suspensions of each organ were centrifuged and resuspended in FACS buffer.

### Sample preparation for senescence quantification

The samples were incubated with LIVE/DEAD Fixable Blue Dead Cell Stain Kit (1:400, Thermo Fisher Scientific, L34961), for 30 minutes at 4°C. Then, the samples were washed with FACS buffer. Next, they were incubated with FcX PLUS anti-mouse CD16/32 (BioLegend, 156604), for 10 minutes at 4°C. Subsequently, the extracellular antibody cocktail prepared in Brilliant Stain Buffer (Thermo Fisher Scientific) was added to the samples (see Supplementary Tables 3-6 for relevant tissue) for 30 minutes at 4°C. The samples were washed and incubated with fixation buffer for 30 minutes at 4°C, before being washed again. A second wash with permeabilization buffer was then performed. The samples were incubated with Donkey Serum (Jackson ImmunoResearch Laboratories) in permeabilization buffer for 5 minutes, before adding the intracellular antibody cocktail (see Supplementary Tables 3-6 for relevant tissue) (30 minutes at 4°C). Of note, we have previously verified the potential of p16 and Bcl-xl as senescence markers^33^, while the potential of γH.2ax was demonstrated following etoposide treatment^34^. Next, the samples were washed three times, once with permeabilization buffer and twice with FACS buffer. Finally, the samples were run using a Cytek Aurora flow cytometer (Cytek Biosciences) and analysed using the SpectroFlo software (Cytek Biosciences), as well as R version 4.4.1 and RStudio version -2024.09.1+394.

### Three-dimensional analysis

Flow cytometry (FCS) data was obtained Cytek Aurora flow cytometer (Cytek Biosciences) and analysed using the SpectroFlo software (Cytek Biosciences). FCS files for each cell type were then exported for subsequent processing. For each distinct cell population identified by these groupings, marker parameter values were log10-transformed and then standardized. Subsequently, the mean and covariance matrix were computed in this transformed space. For visualization purposes, the covariance matrix was then scaled by a constant factor of 0.01 to ensure the resulting ellipsoids were of an appropriate size for clear display, as the large sample sizes (with an average of ∼10,000 cells per sample) would otherwise result in excessively small visual representations. Each processed cell population was then visualized as a 3D ellipsoid, with its centre and shape directly derived from its calculated mean and the scaled covariance matrix. These ellipsoids were rendered together in a single 3D plot, providing a comprehensive visualization of cell population distribution across various cell types and age groups. As an intermediate visual step, a separating manifold was generated for each cell type (within each tissue) to optimally distinguish between the young and old cell populations in the 3D marker space (p16, γH2.ax, Bcl-xl). This manifold was defined by identifying the surface where the probability density of a point belonging to the young population’s distribution equals the probability density of it belonging to the old population’s distribution. Specifically, each population was modelled as a multivariate normal distribution centred at its empirically derived mean. For the purpose of mani fold calculation, a simplified, spherical covariance matix (0.5*IdentityMatrix[3]) was employed to compute the Probability Density Function (PDF) at each point in the 3D space. The separating manifold was then plotted as the ContourPlot3D surface where PDF young(x1,x2,x3) = PDF_old(x1,x2,x3). This manifold was visualized within a 3D plot alongside the respective population ellipsoids. All computational procedures and visualizations were performed using Wolfram Mathematica.

### Heatmap Visualization of Individual Sample Senescence Ranks

A heatmap was generated to visualize the senescence status of individual samples across all analysed cell types and tissues. For each specific cell type within a given tissue, each individual cell event was assigned a quantitative ‘senescence rank.’ This rank was determined by first calculating a projection score for each cell: its marker expression profile (centred by the overall mean of the respective young and old populations for that cell type) was projected onto a vector pointing from this overall mean to the centroid of the old population for that cell type. Subsequently, the ordinal rank of this projection score was assigned to each cell. A higher ordinal rank thus indicated a stronger alignment with the ‘old’ phenotype in that marker space. This approach ensured that the senescence metric was tailored and weighted according to the specific characteristics of each cell type. The heatmap displayed these ranks, with each row representing a unique cell type (from a specific tissue) and columns representing individual samples. All computational procedures and visualizations were performed using Wolfram Mathematica.

### Assessment of marker contributions to the overall variance

To understand the intrinsic contribution of each marker to the overall explained variance, an eigen system analysis, specifically Principal Component Analysis (PCA) on the correlation matrix, was performed for each cell type within each tissue, irrespective of age. This involved calculating the eigenvalues of the correlation matrix, which represent the variance explained by each principal component, and the loadings (eigenvector components), which quantify the contribution of each individual marker (p16, γH2.ax, Bcl-xl) to these components. Results were visualized using pie charts to illustrate the proportion of variance explained by each principal component, and stacked bar charts to show the relative contribution of each marker across all cell types and tissues. All computational procedures and visualizations were performed using Wolfram Mathematica.

### Evaluation of marker combinations for population separation

To assess the necessity of multiple markers for effective population separation, all possible combinations of the 3SMs (p16, γH2.ax, and Bcl-xl) were systematically evaluated. For each cell type within each tissue, a binary classification was performed to distinguish young from old populations. This was achieved by projecting individual cell events onto a derived vector and classifying based on the sign of this projection. The performance of each marker combination was quantified using Balanced Accuracy, calculated as the average of sensitivity and specificity (TP+FNTP+TN+FPTN)/2). This accuracy was computed using the same data from which the population centroids were derived (re-substitution accuracy). The results were then presented in a bar chart illustrating the ‘Fraction of cell types which could be separated with > 75% accuracy’, for each marker combination strategy. All computational procedures and visualizations were performed using Wolfram Mathematica.

### Unbiased quantification of senescent cells

For each cell type within each tissue, a vector representing the direction from the centroid of the young population to the centroid of the old population was calculated. These resultant directional vectors were then normalized and visualized as arrows originating from the origin within a unit sphere. This 3D arrow plot provided insights into age-related shifts in the collective orientation and relative contribution of the p16, γH2.ax, and Bcl-xl markers. For each cell type from a specific tissue, box-whisker plots were generated to visualize the distribution of a derived metric representing the ‘fraction of SnCs’ across different age groups (Young vs. Old). This fraction was determined by first calculating a projection score for each individual cell onto the vector representing the difference between the young and old population centroids for that cell type. A cell was then classified as ‘senescent’ if its projection score was positive (> 0), indicating it falls on the ‘old’ side of the hyperplane orthogonal to the centroid difference vector and passing through the origin in the projected space. The ‘fraction of SnCs’ was then computed as the proportion of cells classified as senescent within each sample. All computational procedures and visualizations were performed using Wolfram Mathematica.

### Opal Multiplex IHC staining

Young and old mice were anaesthetized with Pentobarbital Sodium (CTS Chemical Industries Ltd). Then, mice were perfused with cold PBS for blood clearance. Lungs from these mice were inflated using 4% paraformaldehyde (PFA) before being extracted and fixed in 4% PFA overnight. They were then transferred to 1% PFA for storage, until being embedded in paraffin for histological analysis. Paraffin-embedded tissue sections (4μm) were deparaffinized and rehydrated using Leica Bond dewax solution, followed by blocking of endogenous peroxidase activity with 3% H₂O₂ and 1% HCl in methanol for 30 minutes. Heat-induced antigen retrieval was performed in Tris-EDTA buffer (pH 9.0). To prevent nonspecific binding, sections were incubated with 20% normal horse serum (NHS; VectorLabs, S-2000) and 0.3% Triton X-100. An additional biotin-blocking step (VectorLabs, SP-2001) was incorporated for protocols including biotinylated secondary antibodies. Primary antibodies were diluted in 2% NHS with 0.5% Triton for p16, γH2AX, and Bcl-xl, and in 0.1% Triton for F4/80 and CD45. All staining solutions were freshly prepared, loaded, and processed on the Leica BOND-MAX Fully Automated IHC and ISH Staining System (RRID: SCR_026887) according to the manufacturer’s instructions. The samples were stained with p16 (Abcam, Cat# ab211542, RRID: AB_2891084), Phospho-Histone H2A.X (Ser139) (Cell Signaling Technology, Cat# 2577, RRID: AB_2118010), Bcl-xl (Cell Signaling Technology, Cat# 2764, RRID: AB_2228008), f4/80 (Cell Signaling Technology, Cat# 70076, RRID: AB_2799771) and CD45 (Cell Signaling Technology, Cat# 70257, RRID: AB_2799780). Secondary horseradish peroxidase (HRP)-conjugated antibodies (Jackson ImmunoResearch Labs, Cat# 711-035-152, RRID: AB_10015282) were applied, followed by tyramide signal amplification with fluorescent Opal reagents. Between successive rounds of staining, antibodies were stripped by microwave treatment for 10 min in Tris-EDTA buffer (pH 9.0). The protocol was then repeated from the blocking step.

### Multiplex Fluorescence Imaging

Multispectral imaging was performed using the PhenoImager HT system at 20× magnification (Akoya Biosciences, RRID: SCR_023772) according to the manufacturer’s instructions. The acquired images were subsequently processed using the inForm software (version 3.0; Akoya Biosciences, RRID: SCR_019155) for spectral unmixing and background subtraction. To correct for tissue autofluorescence, an unstained slide (without DAPI or Opal fluorophores) was used, and regions exhibiting high autofluorescence were annotated within inForm for processing. Image stitching was performed automatically by the PhenoImager HT software.

### Single-cell RNA sequencing

The lungs from 24-months old mice were collected and dissociated following the method described above (in the ‘Organs dissociation to a single-cell suspension’ section). The cells collected were then incubated with a Zombie NIR viability dye (BioLegend, 423106) for 30 minutes at 4°C. Next, the samples were labelled and fixed following the 10X Genomics protocol: ‘Cell surface and intracellular Protein labelling for Chromium fixed RNA profiling’ (10xgenomics, CG000529). The samples were stained with the extracellular marker CD45 Pacific Blue (1:100, BioLegend, 103126). Later, they were stained with the intracellular markers anti-p16 (1:200, Abcam, ab54210) conjugated to Alexa Fluor 647 (Thermo Fisher Scientific, A-20186), anti-γH2.ax PE (1:100, BD Biosciences, 562377) and anti-Bcl-xl PE-Cy7 (1:200, Cell Signaling Technology, 81965). These cells were sorted for immune (CD45+) and non-immune (CD45-) SnCs (triple positive for the 3SMs), using the BD FACSDiscover S8 Cell Sorter and frozen for later processing. Single cell RNA-seq libraries were prepared using the Chromium fixed RNA profiling for multiplexed samples according to the manufacturer’s protocol using Chromium Fixed RNA Kit, Mouse Transcriptome (10x genomics). Fixed samples were multiplexed and loaded onto Next GEM Chip Q targeting 10,000 cells per sample and then ran on a Chromium Controller instrument to generate GEM emulsion (10x Genomics), followed by library preparation. Final libraries were quantified using NEBNext Library Quant Kit for Illumina (NEB) and high sensitivity D1000 TapeStation (Agilent). Libraries were sequenced on a NovaSeq X Plus instrument using a 1.5B 100 cycles reagent kit (Illumina).

### scRNAseq analysis

Single cell RNA sequences (scRNAseq) data files were acquired from Cell Ranger v8 (“10x Genomics”) and subjected to default parameters for alignment, filtering, barcode counting, and UMI counting. Total lung scRNAseq data was obtained from Gene Expression Omnibus (GEO, deposited as GSE141259)^24^. Initial quality control measurements were performed as described by Strunz et al.^24^. For comparative purposes, cells from ‘whole lung’ dataset of control mice injected with PBS on day 0 were used. Coherently, the samples’ cell types were annotated using the same annotations^24^ recovered from ‘metacelltype’ metadata column. Rare un-matched cells annotations were removed. Data manipulation was carried using Seurat v5 platform in R 4.4.0 and R studio 2025.05.0+496. Cells with less than 200 features, below or above 4 Median Absolute Deviations (MAD) of nUMIs, nCounts and percents of mitochondrial genes were removed. Also, genes found in less than 5 cells were removed. Then, objects were normalized using ‘SCTransform’, and samples were integrated using “PrepSCTIntegration”, “FindIntegrationAnchors” and “IntegrateData”, as described by Stuart et al.^35^. Afterwards, the data was split into 2 objects, resident and immune cells and all analyses were performed separately. Objects were re-normalized using ‘SCTransform’, and ‘RunPCA’ for data reduction and preparation for layers integration using ‘IntegrateLayers’; followed by ‘FindNeighbors’, ‘FindClusters’ and ‘RunUMAP’ for presentation of cells’ coordinates on top of dimension plots. Seurat objects were converted into AnnData compatible format (“.h5ad”) for ‘SenePy’ scoring^25^ in Python 3.11.7 Jupyter Notebook 7.3.2. Specific cell types scoring was assigned when relevant matching cell type annotations were applicable. SenePy scores were transitioned into corresponding Seurat objects for creation of violin plots. The UMAPs representing the SenePy score color scale were created using the log1p(senepy_score) function in R, using the natural base e (∼2.718). The function adds 1 to the senepy_score to avoid impossible log(zero), and the equation is as follows: loge(senepy_score + 1).

### Senescence quantification assay of human PBMCs

Frozen PBMC samples from young (>35y.o.) and old donors (<65y.o.) were purchased (Charles River Laboratories) for this experiment. These PBMCs were thawed in RPMI supplemented with 10% FBS, 100 units/ml of penicillin and 100 mg/ml of streptomycin. They were then centrifuged and resuspended in PBS, where they were incubated with LIVE/DEAD Fixable Blue Dead Cell Stain Kit (1:400, Thermo Fisher Scientific, L34961), for 30 minutes at 4°C. The samples were then washed and incubated in Brilliant Stain Buffer (Thermo Fisher Scientific) with the extracellular antibody cocktail (see Supplementary Table 7) for 30 minutes at 4°C. Next, they were washed and incubated with fixation buffer for 30 minutes at 4°C. The samples were washed twice, once with FACS buffer and once with permeabilization buffer. The samples were incubated with Donkey Serum (Jackson ImmunoResearch Laboratories) in permeabilization buffer for 5 minutes, and stained with the 3SMs, anti-p16 AF647 (1:1000, Abcam, ab192054), anti-γH2.ax PE (1:1000, BD Biosciences, 562377) and anti-Bcl-xl PE-Cy7 (1:1000, Cell Signaling Technology, 81965) (30 minutes at 4°C). Then, the samples were washed once with permeabilization buffer and twice with FACS buffer. Finally, the samples were run using a Cytek Aurora flow cytometer (Cytek Biosciences) and analysed using the SpectroFlo software (Cytek Biosciences).

### Heatmap

Heatmaps were obtained using R version 4.4.1 and RStudio version -2024.09.1+394, using the “ggplot2” library. For the creation of the heatmap in Extended Data Fig. 2c, the relative means of old vs. young cell’s composition were calculated in each of the corresponding organs’ cell types. The heatmap in Extended Data Fig. 5b was created similarly but represents the relative means of old vs. young SnCs’ percentages.

### Correlation analysis

R version 4.4.1 and RStudio version -2024.09.1+394 were used to create the correlation plots, using the “corrplot” library. “Pearson” correlation of old mice percentages of senescent blood cells vs. - senescent liver, - senescent lung and - senescent intestine cells was calculated. To enforce adequate computation, only non-missing values in both organs’ columns were used for correlation calculation.

### Statistical analysis

The data acquired is presented in the figures as mean values ±SEM. Statistical analysis of the senescence quantification using manual gating was performed using the GraphPad Prism software (GraphPad Software), and determined using one sample t-tests. A P value < 0.05 was considered statistically significant (* < 0.05, ** < 0.01, *** < 0.005). Statistical analysis of the unbiased senescence quantification was performed using a Mann-Whitney U test. The p-values from these tests were then adjusted for multiple comparisons using the Bonferroni correction, with significance levels indicated on the plots: * for p < 0.05, ** for p < 0.001, and *** for p < 0.0001 (after Bonferroni adjustment). Statistical analysis of the scRNAseq SenePy score results was performed using a Wilcoxon test to generate p-values, using the ‘compare_means’ function of the ggpubr package in R. A P value < 0.05 was considered statistically significant (* < 0.05, ** < 0.01, *** < 0.001, **** < 0.0001).

## Acknowledgments

We sincerely thank Papismadov N. for helpful advice and comments on the manuscript. We also thank all members of the Krizhanovsky laboratory for helpful discussions. V.K. was supported by grants from the European Research Council (856487), from the Israel Science Foundation (1626/20), DFG - CRC 1506 “Aging at Interfaces”, Weizmann - Nella and Leon Benoziyo Center for Neurological Diseases, Weizmann - Sagol Center for Research on the Aging Brain, Weizmann SABRA - Yeda-Sela - WRC Program, the Estate of Emile Mimran, and The Maurice and Vivienne Wohl Biology Endowment, the Estate of Gerald Alexander and Moross Integrated Cancer Center V.K. is the Director of EKARD Institute for Cancer Diagnosis Research.

## Contributions

VK, HG and UC conceived the outline of the project and conceptualized it. UC, HG, IS and VK planned the experiments. UC performed and analysed the *in vitro* Flow Cytometry experiments. UC, HG and IS designed and performed the mice Flow Cytometry experiments. UC, HG, IS and EK established the antibody panels and analysed the mice Flow Cytometry Data (initial analysis and manual gating). AM performed the three-dimensional analysis, the analysis of the contribution of each marker and the PCA analysis. HA and NR helped with the mice sacrifices and organ recovery. OM performed and analysed the immunohistochemistry experiments. UC, EK, RB and HKS performed the scRNAseq experiments. HA analysed the scRNAseq data. UC, IS and HA performed the correlation analysis. UC performed and analysed the hPBMCs Flow Cytometry experiments. UC and VK wrote the manuscript. VK was responsible for funding acquisition and supervised the study. All authors read and approved the final manuscript.

