## Extended Data Figures and Tables for "Single-cell quantification of senescence burden reveals cell type-specific ageing dynamics across organs"

### Figure legends

**Extended Data Figure 1: The percentage of different cell types as it was recovered from the lung, liver, intestine and blood of young and old mice. (a)** Pie charts representing the proportion of immune or resident cells in young and old mice, in lung, liver and intestine tissues, and blood. **(b)** Heatmap representing the log10 ratio of the proportion of each cell type in old over young mice. Each line represents a different organ as indicated on the left. Epi- Epithelial cells; End- endothelial cells; Fib- fibroblasts; Mac- macrophages; Mon- monocytes; Neu- Neutrophils; DC- dendritic cells; NK- NK cells; B- B cells; CD4- CD4 T cells; CD8- CD8 T cells; AM- alveolar macrophages; IM- Interstitial macrophages.

**Extended Data Figure 2: Combination of senescence markers distinguishes between young and old mice in multiple organs and cell types.** A three-dimensional analysis using the 3SMs, helps define a 'new' composite senescence score based on the expression of these markers, and define a plane which can separate young from old mice in each cell type. This analysis automatically finds the best separating manifold by using a support vector machine (SVM) to define a separating plane and a separating direction between young and old groups. **(a)** lungs **(b)** liver **(c)** intestine **(d)** blood.

**Extended Data Figure 3: Senescent macrophages can be identified in the lung of old mice by immunostaining**. Multispectral imaging of lung tissues from young and old mice using the PhenoImager HT system with antibodies directed to CD45 and F4/80 to identify macrophages and antibodies directed to our selected senescence markers, γH.2ax, p16 and Bcl-xl. Arrows indicate cells positive for all markers. Scale bars, 20μm and 2μm.

**Extended Data Figure 4: SenM+ cells quantification based on the 3-dimensional analysis, shows significant increases in senescence burden in lung and intestinal immune populations. (a)** Quantification of the percentage of blood circulating endothelial cells positive for the selected senescence markers (SenM+). End- endothelial cells. **(b)** Heatmap representing the log10 ratio of the proportion of SenM+ cells in old over young mice. Each line represents a different organ as indicated on the left. Epi- Epithelial cells; End- endothelial cells; Fib- fibroblasts; Mac- macrophages; Mon- monocytes; Neu- Neutrophils; DC- dendritic cells; NK- NK cells; B- B cells; CD4- CD4 T cells; CD8- CD8 T cells. **(c)** The arrows represent the direction in which the expression of senescence markers evolves with age. Only cell types which moved in the direction of the blue arrows were selected (all 3SMs expression increases with age). **(d)** Quantification of SenM+ populations in relevant cell types in the lung and intestine of young and old mice was obtained in an unbiased manner, by applying a threshold on the expression distribution of the selected senescent markers, above which cells were considered SenM+. A P value $< 0.05$ was considered statistically significant, following a Mann-Whitney U test (* < 0.05, ** < 0.001, *** < 0.0001 (after Bonferroni adjustment)). CD8- CD8 T cells; NK- NK cells; B- B cells; AM- alveolar macrophages; IM- interstitial macrophages; Ly6C- -Ly6C- monocytes; Mon- monocytes; DC- dendritic cells; Mac- macrophages.

**Extended Data Figure 5: SenePy score comparisons of SenM+ cell types to matched total lung cell types using cell type specific SenePy signatures. (a)** UMAP of SenM+;CD45- and total lung CD45- cells labelled by cell types. **(b) Picked cell type from Fig. 4e.** SenM+ and total lung fibroblasts violin plots of SenePy scores based on the cell-type-specific SenePy signature. **(c)** UMAP of SenM+;CD45+ and total lung CD45+ cells labelled by broad cell types. **(d) Picked cell types from Fig. 4f.** SenM+ and total lung DCs and granulocytes violin plots of SenePy scores based on the cell-type-specific SenePy signature. **(e) Picked cell types from Fig. 4f** SenM+ and total lung macrophages and monocytes violin plots of SenePy scores based on the cell-type-specific SenePy signature. **(f) Picked cell type from Fig. 4f** SenM+ and total lung B, NK and T cells violin plots of SenePy scores based on the cell-type-specific SenePy signature A P value < 0.05 was considered statistically significant, following a Wilcoxon test analysis (* < 0.05, ** < 0.01, *** < 0.001, **** < 0.0001).

**Extended Data Figure 6: Cumulative senescence burden of each mouse per tissue. (a)** liver **(b)** intestine **(c)** Blood. The cumulative senescence burden of each mouse for each organ was calculated by adding together the percentages of senescent cells in each of their cell types. Each bar represents a single mouse.

**Extended Data Figure 7: Correlation analysis of SenM+ cell burden between different cell types in (a) liver (b) intestine and blood of aged mice.** A correlation plot comparing each cell types in the liver and blood or intestine and blood of aged mice, was created using R and RStudio with "corrplot" libraries.

**Extended Data Figure 8: Correlation analysis of SenM+ cell burden between different cell types in (a) lung and liver (b) liver and intestine (c) lung and intestine of aged mice.** A correlation plot comparing each cell types in the lung and liver, liver and intestine or lung and intestine of aged mice, was created using R and RStudio with "corrplot" libraries.

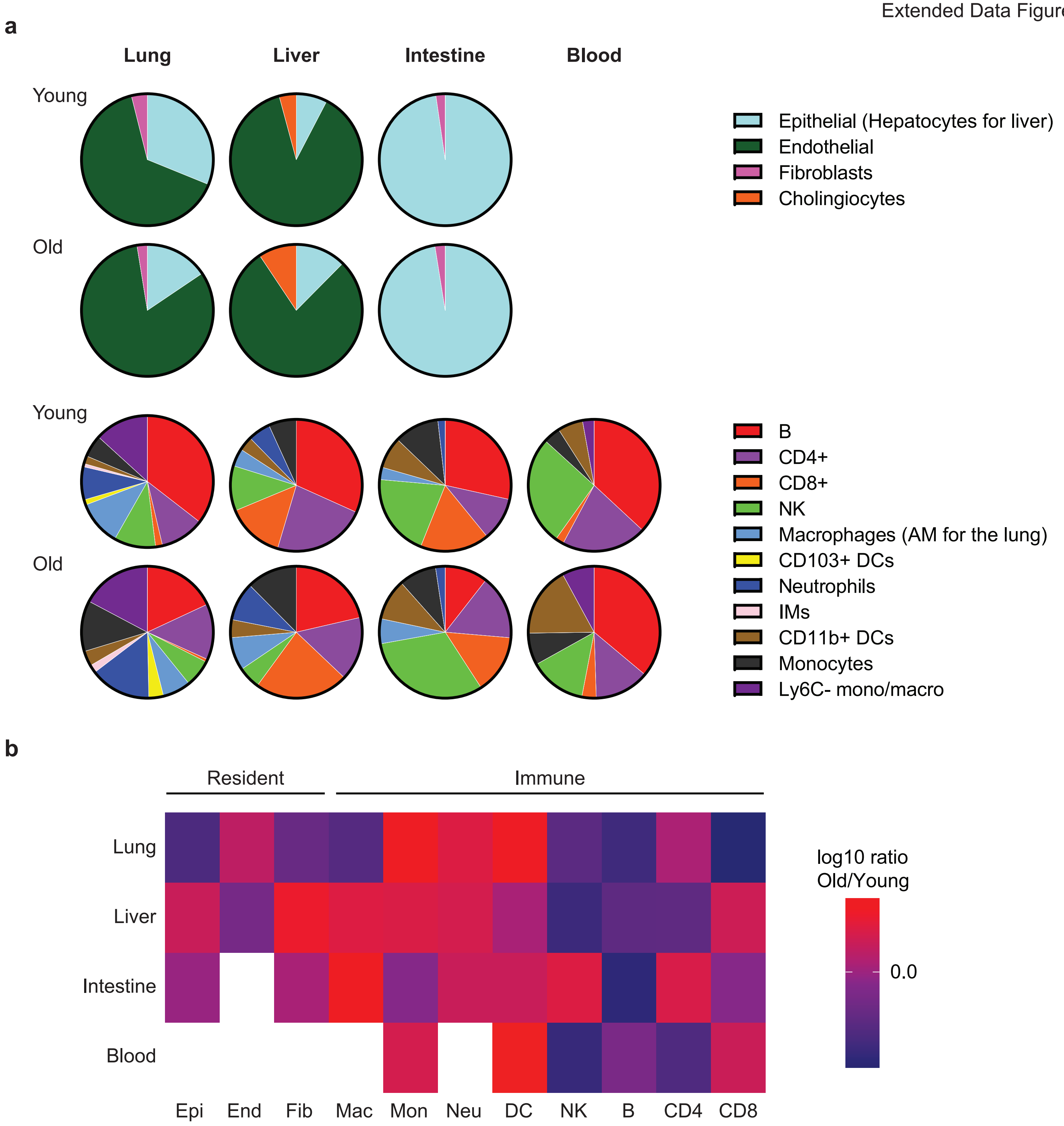
a
Lung
Liver
Intestine
Blood
Young
Old
Young
Old
Epithelial (Hepatocytes for liver)
Endothelial
Fibroblasts
Cholingiocytes
B
CD4+
CD8+
NK
Macrophages (AM for the lung)
CD103+ DCs
Neutrophils
IMs
CD11b+ DCs
Monocytes
Ly6C- mono/macro
b
Resident
Immune
Lung
Liver
Intestine
Blood
Epi
End
Fib
Mac
Mon
Neu
DC
NK
B
CD4
CD8
log10 ratio
Old/Young
0.0

Extended Data Figure 2

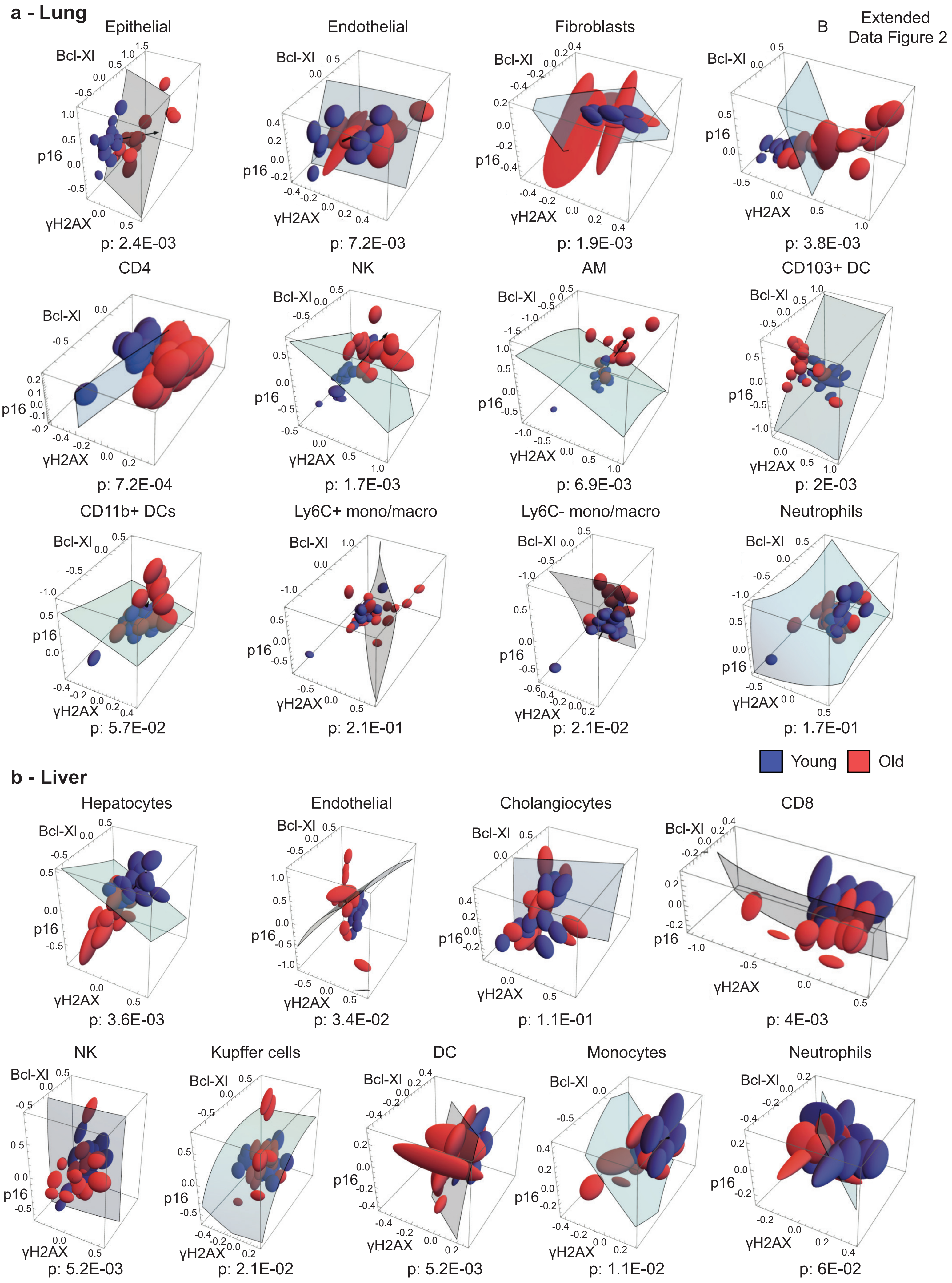

**c Intestine**

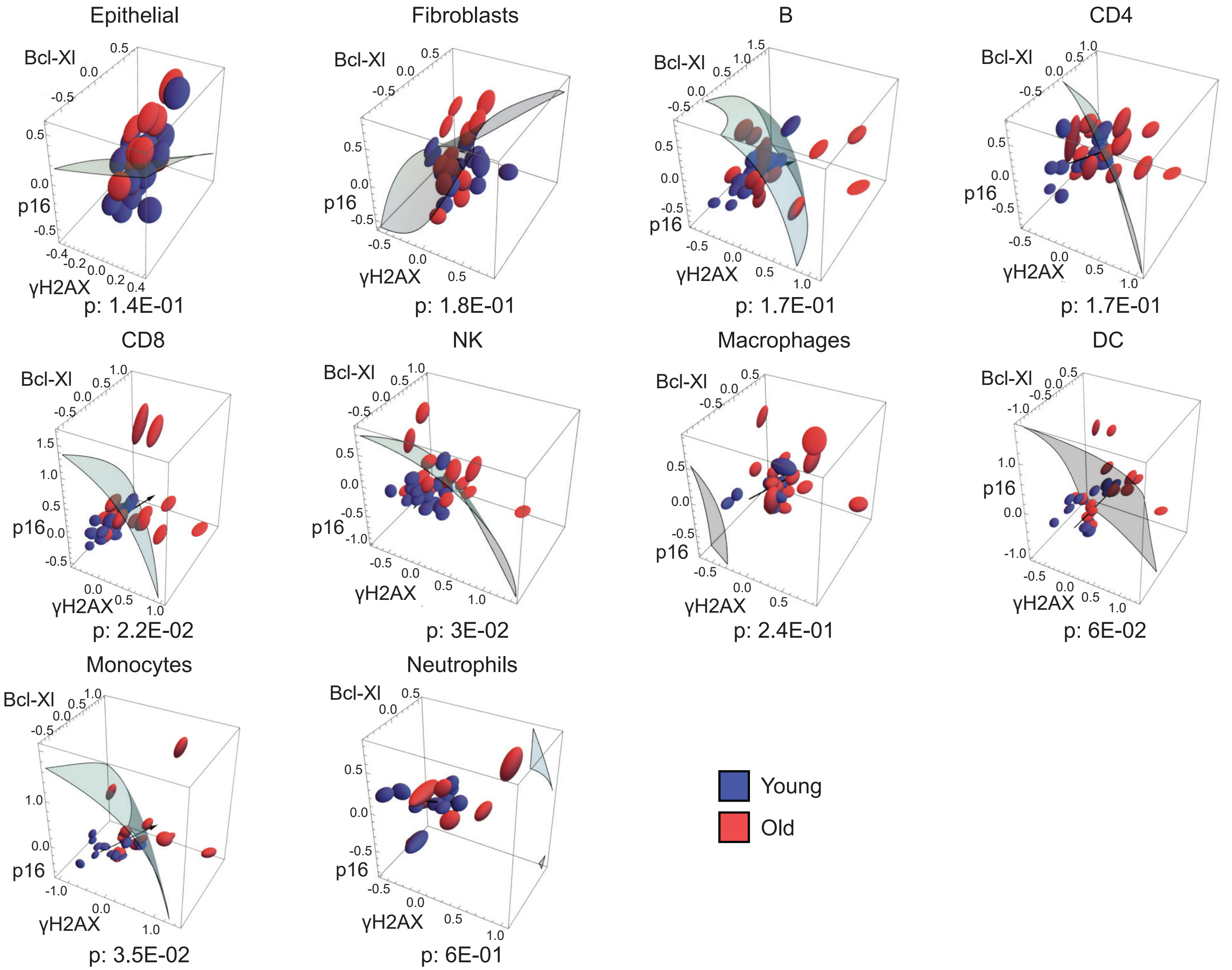


**d Blood**

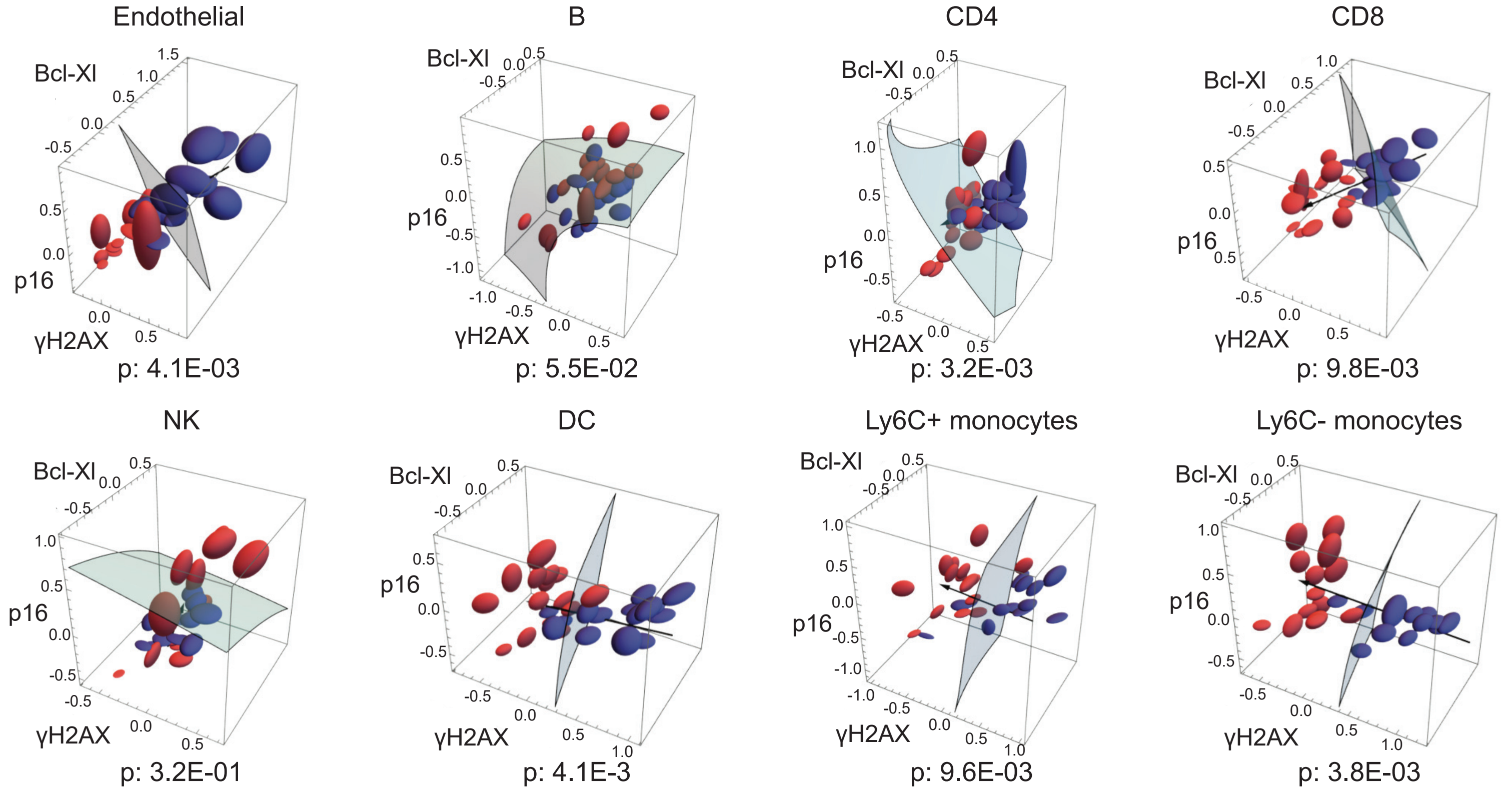

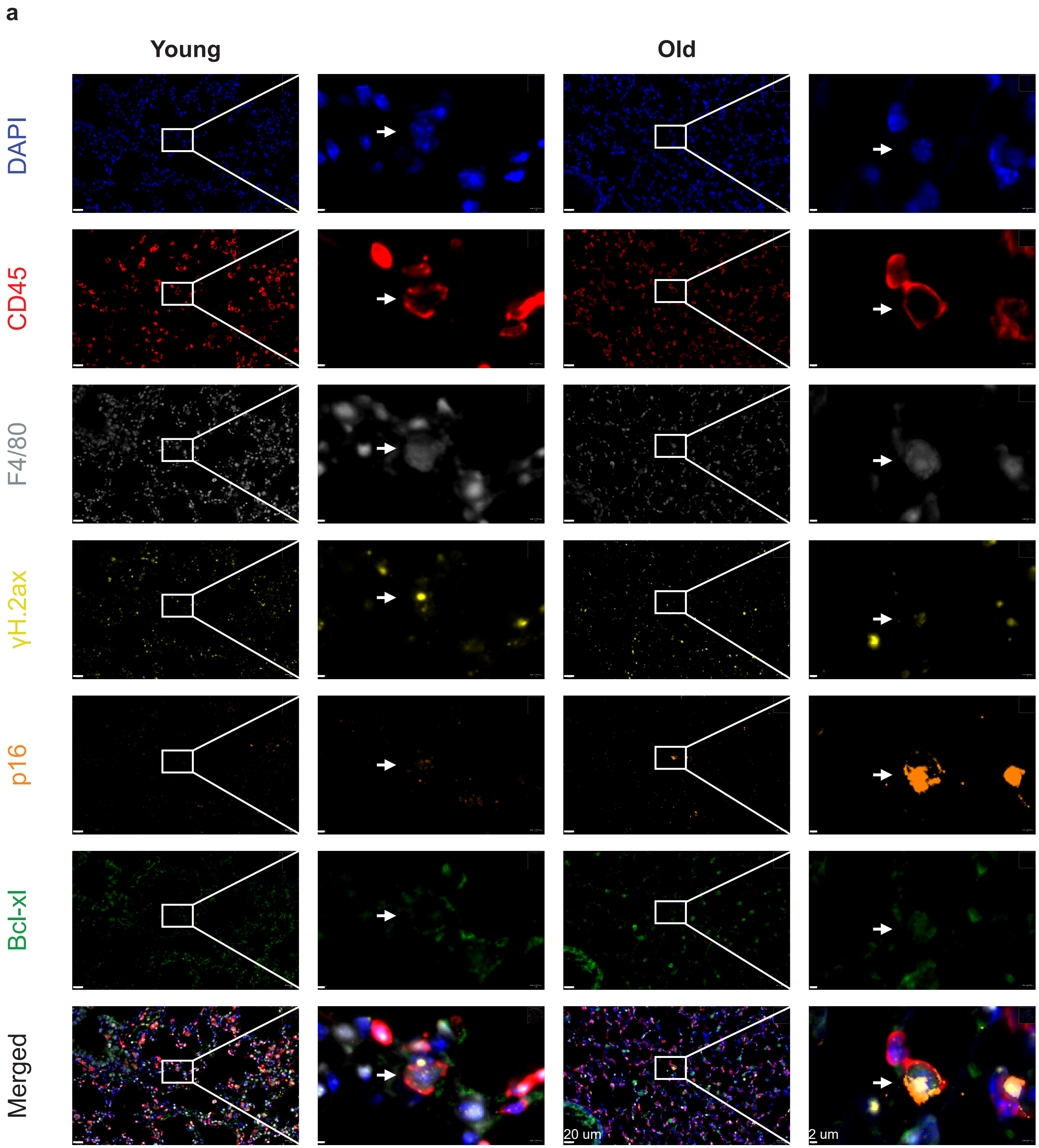
a
Young
Old
DAPI
CD45
F4/80
γH.2ax
p16
Bcl-xl
Merged
20 um
2 um

Extended Data Figure 4

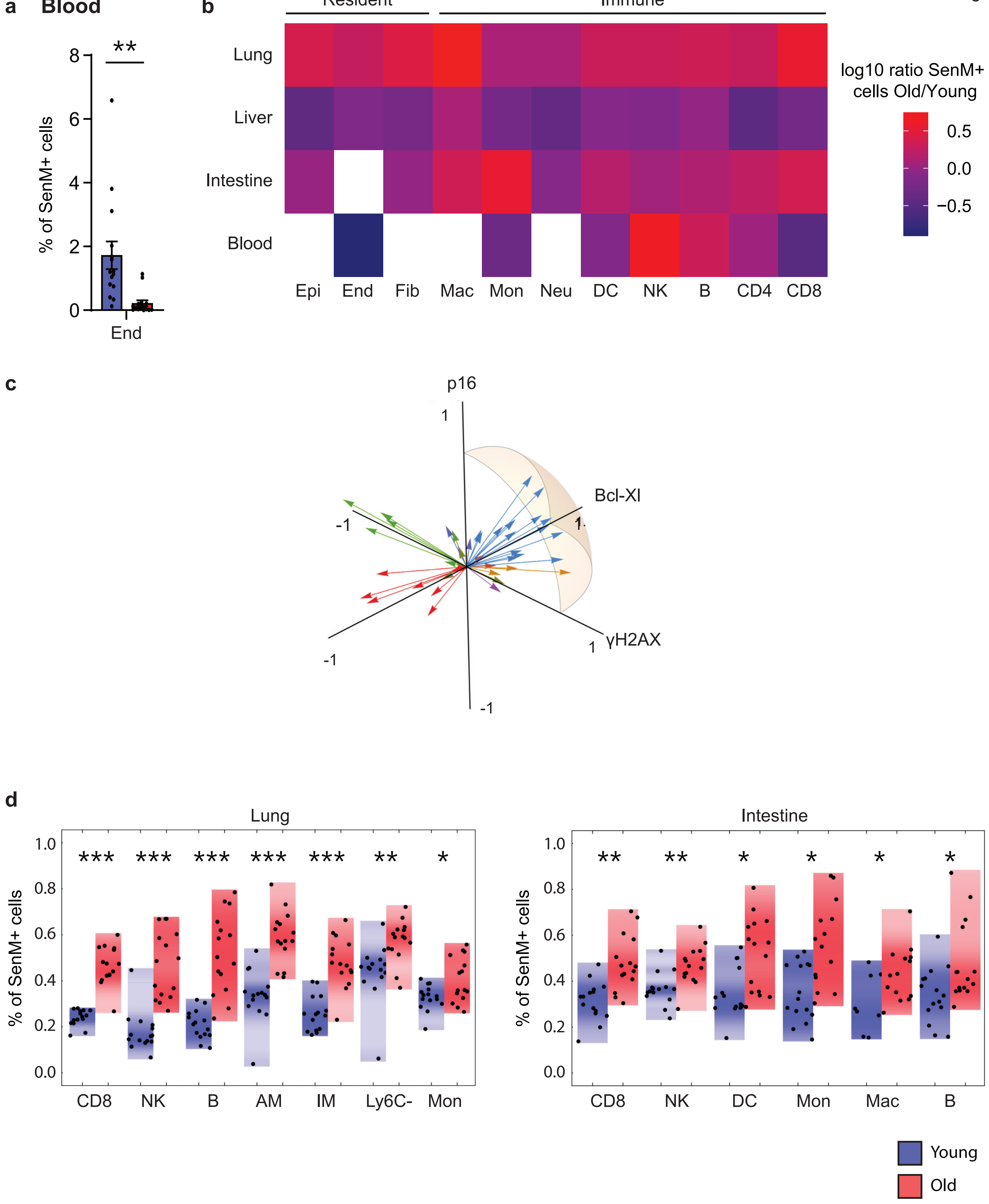

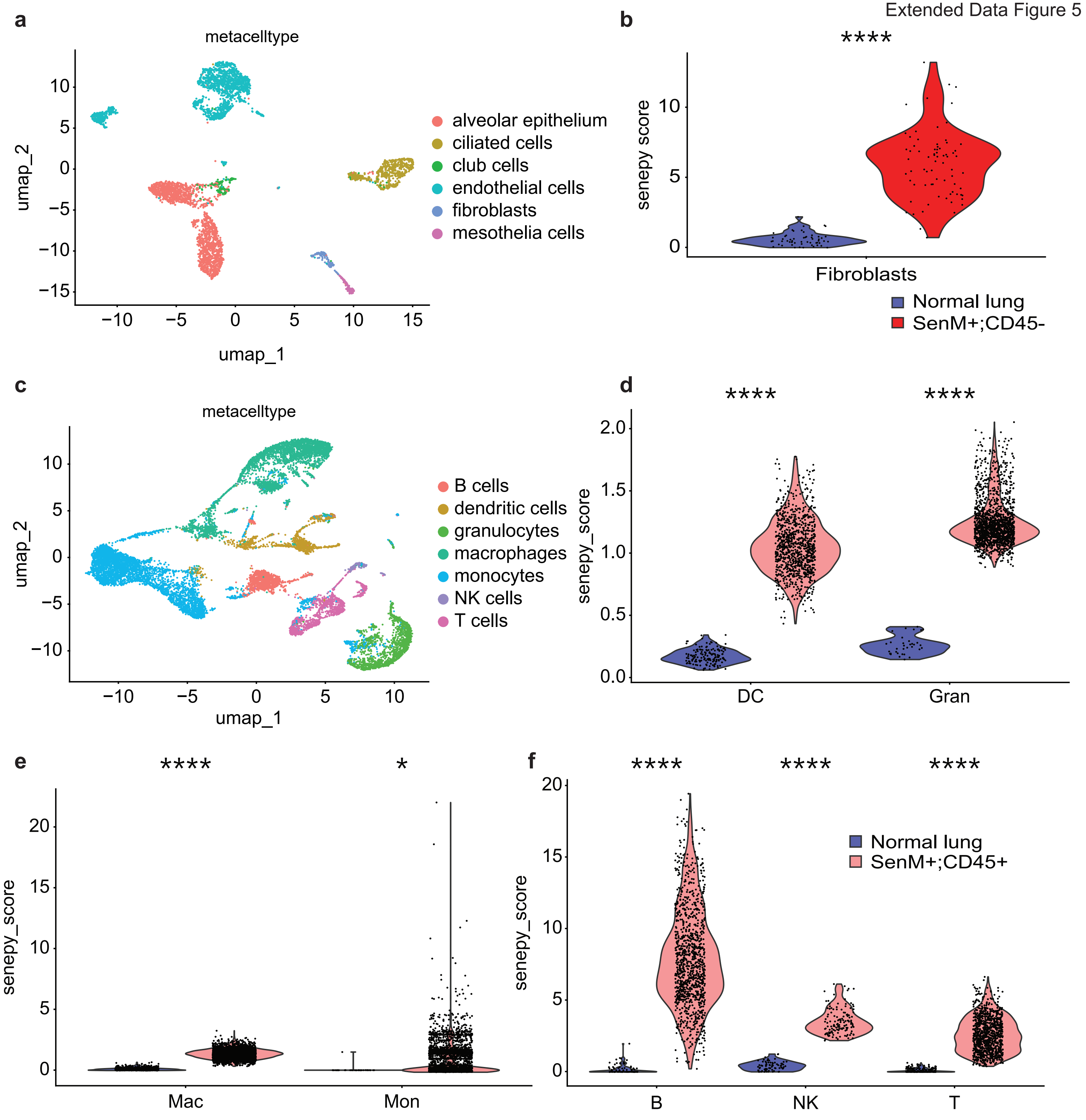
a
metacelltype
umap_2
umap_1
alveolar epithelium
ciliated cells
club cells
endothelial cells
fibroblasts
mesothelia cells
b
****
senepy score
Fibroblasts
Normal lung
SenM+;CD45-
c
metacelltype
umap_2
umap_1
B cells
dendritic cells
granulocytes
macrophages
monocytes
NK cells
T cells
d
****
****
senepy_score
DC
Gran
e
****
*
senepy_score
Mac
Mon
f
****
****
****
senepy_score
Normal lung
SenM+;CD45+
B
NK
T

**a**

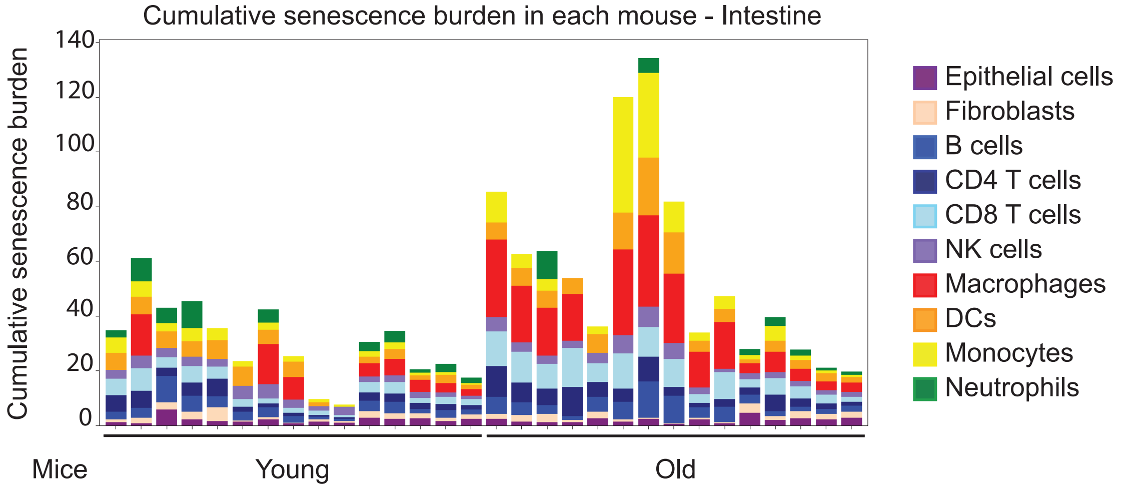
Cumulative senescence burden in each mouse - Intestine
Cumulative senescence burden
140
120
100
80
60
40
20
0
Mice
Young
Old
Epithelial cells
Fibroblasts
B cells
CD4 T cells
CD8 T cells
NK cells
Macrophages
DCs
Monocytes
Neutrophils

**b**

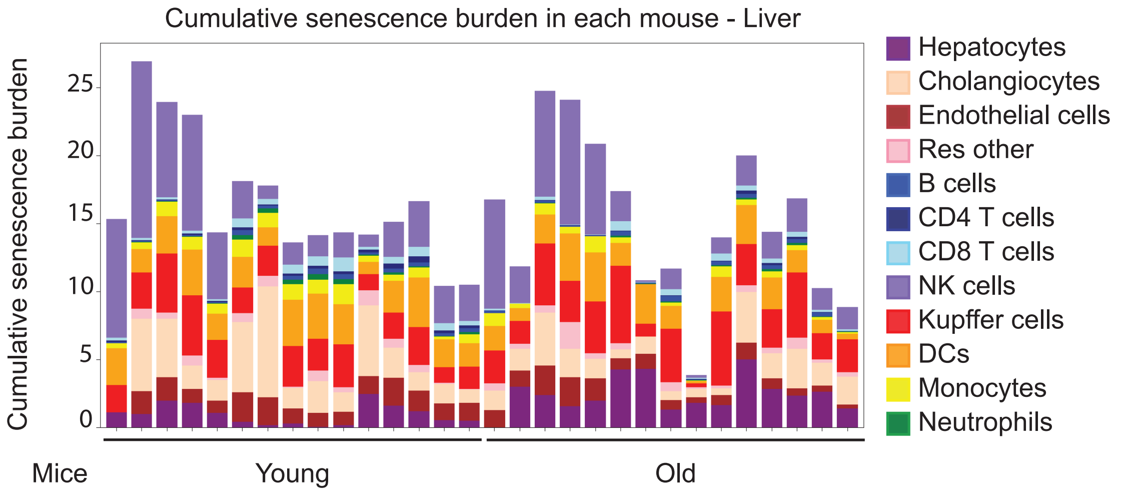
Cumulative senescence burden in each mouse - Liver
Cumulative senescence burden
25
20
15
10
5
0
Mice
Young
Old
Hepatocytes
Cholangiocytes
Endothelial cells
Res other
B cells
CD4 T cells
CD8 T cells
NK cells
Kupffer cells
DCs
Monocytes
Neutrophils

**c**

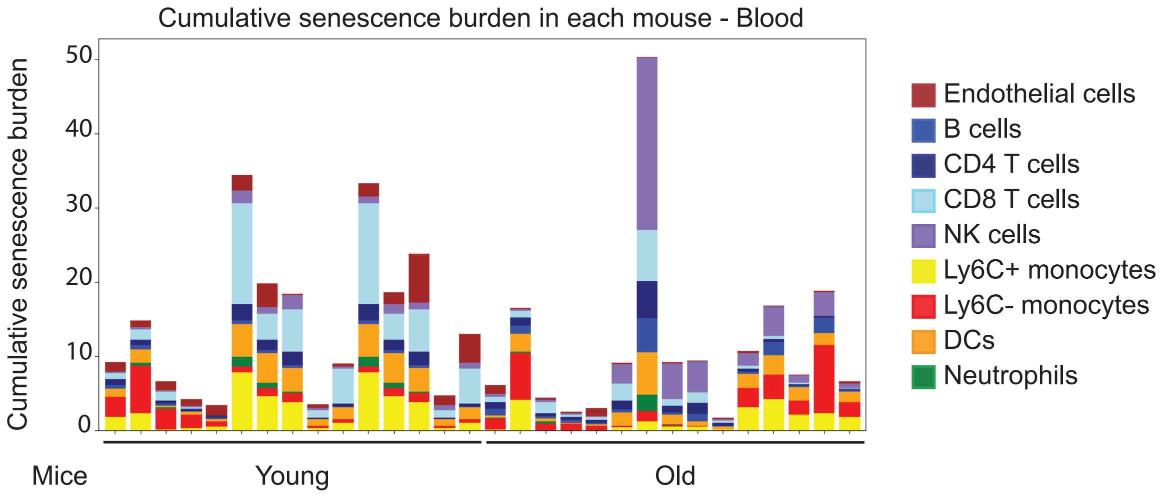
Cumulative senescence burden in each mouse - Blood
Cumulative senescence burden
50
40
30
20
10
0
Mice
Young
Old
Endothelial cells
B cells
CD4 T cells
CD8 T cells
NK cells
Ly6C+ monocytes
Ly6C- monocytes
DCs
Neutrophils

**a**

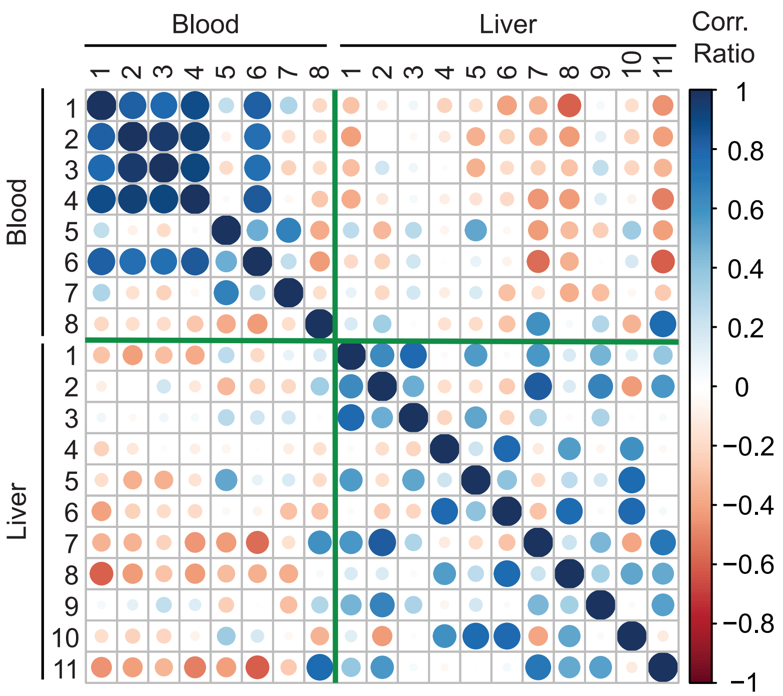


Blood

1. B cells
2. CD4 T cells
3. CD8 T cells
4. NK cells
5. Ly6C+ mono/macro
6. DCs
7. Ly6C- mono/macro
8. Endothelial cells

Liver

1. Hepatocytes
2. Endothelial cells
3. Cholangiocytes
4. B cells
5. CD4 T cells
6. CD8 T cells
7. NK cells
8. Kupffer cells
9. Dendritic cells
10. Neutrophils
11. Monocytes

**b**

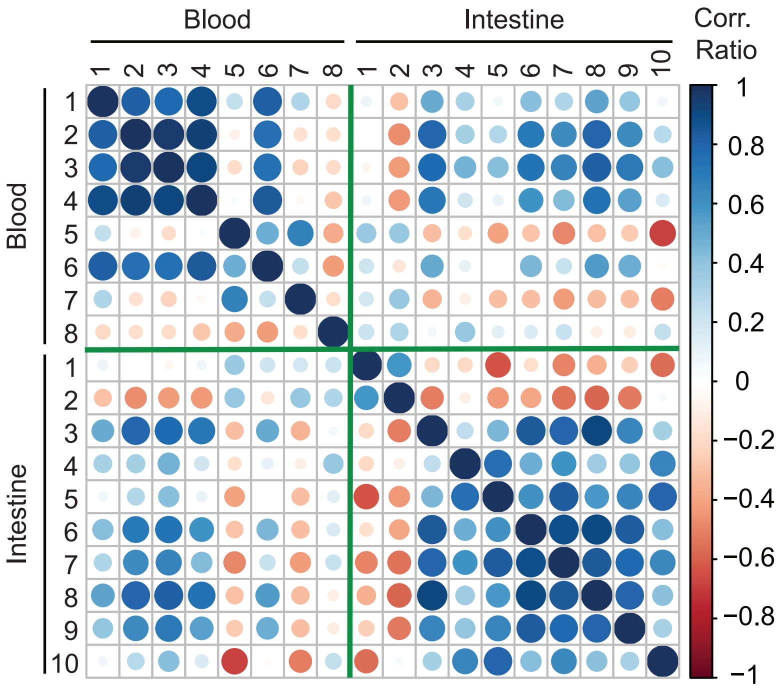


Blood

1. B cells
2. CD4 T cells
3. CD8 T cells
4. NK cells
5. Ly6C+ mono/macro
6. DCs
7. Ly6C- mono/macro
8. Endothelial cells

Intestine

1. Epithelial cells
2. Fibroblasts
3. B cells
4. CD4 T cells
5. CD8 T cells
6. NK cells
7. Macrophages
8. Dendritic cells
9. Monocytes
10. Neutrophils

Extended Data Figure 8

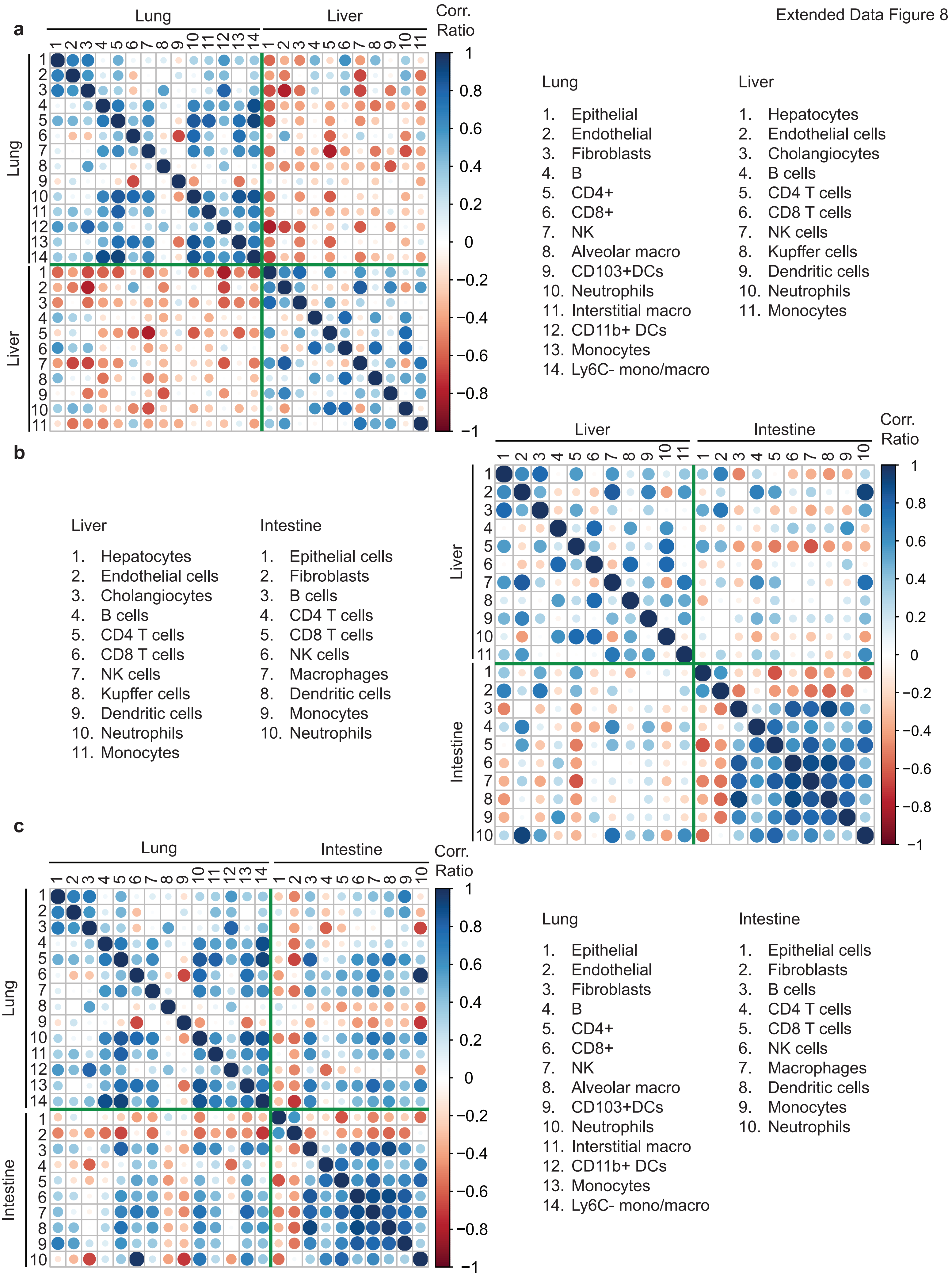

| Significant separation between young and old | | |
|---|---|---|
| **Tissue** | **Cell type** | **P-value** |
| Lung | Epithelial cells | 0.0024 |
| Lung | Endothelial cells | 0.0072 |
| Lung | Fibroblasts | 0.0019 |
| Lung | B cells | 0.0038 |
| Lung | CD4 T cells | 0.0007 |
| Lung | CD8 T cells | 0.0015 |
| Lung | NK cells | 0.0017 |
| Lung | Alveolar macrophages | 0.0069 |
| Lung | Interstitial macrophages | 0.002 |
| Lung | CD103+ DCs | 0.002 |
| Lung | Ly6C- monocytes | 0.0205 |
| Liver | Hepatocytes | 0.0036 |
| Liver | Endothelial cells | 0.0344 |
| Liver | B cells | 0.0029 |
| Liver | CD4 T cells | 0.0008 |
| Liver | CD8 T cells | 0.004 |
| Liver | NK cells | 0.0052 |
| Liver | Kupffer cells | 0.0211 |
| Liver | DCs | 0.0052 |
| Liver | Monocytes | 0.0106 |
| Intestine | CD8 T cells | 0.0225 |
| Intestine | NK cells | 0.0305 |
| Intestine | Monocytes | 0.035 |
| Blood | Endothelial cells | 0.0041 |
| Blood | CD4 T cells | 0.0032 |
| Blood | CD8 T cells | 0.0098 |
| Blood | DCs | 0.0041 |
| Blood | Ly6C+ monocytes | 0.0096 |
| Blood | Ly6C- monocytes | 0.0038 |

**Supplementary table 1: Cell types showing significant separations between young and old groups based on 3-dimensional analysis using three senescence markers (p16, γH.2ax and Bcl-xl).**

| No significant separation between young and old | | |
|---|---|---|
| **Tissue** | **Cell type** | **P-value** |
| Lung | CD11b+ DCs | 0.0568 |
| Lung | Ly6C+ monocytes | 0.2053 |
| Lung | Neutrophils | 0.1745 |
| Liver | Cholangiocytes | 0.1106 |
| Liver | Neutrophils | 0.0597 |
| Intestine | Epithelial cells | 0.1392 |
| Intestine | Fibroblasts | 0.1847 |
| Intestine | CD4 T cells | 0.1671 |
| Intestine | B cells | 0.173 |
| Intestine | Macrophages | 0.2382 |
| Intestine | DCs | 0.0597 |
| Intestine | Neutrophils | 0.5986 |
| Blood | B cells | 0.0552 |
| Blood | NK cells | 0.3228 |

**Supplementary table 2: Cell types which do not show significant separations between young and old groups based on 3-dimensional analysis using three senescence markers (p16, γH.2ax and Bcl-xl).**

| Lung | | |
|---|---|---|
| Extracellular antibody panel | | |
| **Antibody dilution** | **Clone** | **Cat. No.** |
| **Non-immune panel** | | |
| BV421 CD140a (1:100) | APA5 | 135923, BioLegend |
| BUV395 Ep-CAM (1:100) | G8.8 | 740281, BD Biosciences |
| BV605 CD31 (1:200) | 390 | 102427, BioLegend |
| BUV496 CD45 (1:100) | 30-F11 | 749889, BD Biosciences |
| **Immune panel** | | |
| BUV496 CD45 (1:100) | 30-F11 | 749889, BD Biosciences |
| AF488 CD45R/B220 (1:100) | RA3-6B2 | 103225, BioLegend |
| AF532 Ly-6C (1:100) | ER-MP20 | NB100-65413AF532, Novus |
| BUV737 Ly-6G (1:100) | 1A8 | 741813, BD Biosciences |
| BV510 CD11b (1:100) | M1/70 | 101245, BioLegend |
| BV711 CD64 (1:100) | X54-5/7.1 | 139311, BioLegend |
| BV650 CD3 (1:100) | 17A2 | 100229, BioLegend |
| BV570 CD4 (1:100) | RM4-5 | 100541, BioLegend |
| PerCP-eFluor 710 CD8a (1:100) | 53-6.7 | 46-0081-82, Thermo Fisher Scientific |
| Vioblue CD11c (1:100) | N418 | 130-102-797, Miltenyi Biotec |
| BV785 MHCII (1:250) | M5/114.15.2 | 107645, BioLegend |
| AF700 NK-1.1 (1:100) | PK136 | 108729, BioLegend |
| **Intracellular antibody panel** | | |
| PE-Cy7 Bcl-xl (1:200) | 54H6 | 81965, Cell Signaling Technology |
| PE H2AX (1:100) | N1-431 | 562377, BD Biosciences |
| Purified recombinant p16 (1:200) (conjugated to AF647) | 2D9A12 | ab54210, Abcam |
| Alexa Fluor™ Antibody Labeling Kits | | A-20186, Thermo Fisher Scientific |

**Supplementary table 3: Extracellular and Intracellular antibody panels used to stain mice lungs**

| Liver | | |
|---|---|---|
| **Extracellular antibody panel** | | |
| **Antibody dilution** | **Clone** | **Cat. No.** |
| **Non-immune panel** | | |
| FITC CD26 (1:100) | H194-112 | 137805, BioLegend |
| BV605 CD31 (1:200) | 390 | 102427, BioLegend |
| BUV496 CD45 (1:100) | 30-F11 | 749889, BD Biosciences |
| **Immune panel** | | |
| BUV496 CD45 (1:100) | 30-F11 | 749889, BD Biosciences |
| AF488 CD45R/B220 (1:100) | RA3-6B2 | 103225, BioLegend |
| AF532 Ly-6C (1:100) | ER-MP20 | NB100-65413AF532, Novus |
| BUV737 Ly-6G (1:100) | 1A8 | 741813, BD Biosciences |
| BV510 CD11b (1:100) | M1/70 | 101245, BioLegend |
| BV711 CD64 (1:100) | X54-5/7.1 | 139311, BioLegend |
| BV650 CD3 (1:100) | 17A2 | 100229, BioLegend |
| BV570 CD4 (1:100) | RM4-5 | 100541, BioLegend |
| PerCP-eFluor 710 CD8a (1:100) | 53-6.7 | 46-0081-82, Thermo Fisher Scientific |
| Vioblue CD11c (1:100) | N418 | 130-102-797, Miltenyi Biotec |
| BV785 MHCII (1:250) | M5/114.15.2 | 107645, BioLegend |
| AF700 NK-1.1 (1:100) | PK136 | 108729, BioLegend |
| APC-Cy7 F4/80 (1:100) | BM8 | 123117, BioLegend |
| **Intracellular antibody panel** | | |
| AF594 Cytokeratin19 (1:100) (non-immune panel) | EP1580Y | ab203443, Abcam |
| PE-Cy7 Bcl-xl (1:200) | 54H6 | 81965, Cell Signaling Technology |
| PE H2AX (1:200) | N1-431 | 562377, BD Biosciences |
| Purified recombinant p16 (1:200) (conjugated to AF647) | 2D9A12 | ab54210, Abcam |
| Alexa Fluor™ Antibody Labeling Kits | | A-20186, Thermo Fisher Scientific |

**Supplementary table 4: Extracellular and Intracellular antibody panels used to stain mice livers**

| Intestine | | |
|---|---|---|
| **Extracellular antibody panel** | | |
| **Antibody dilution** | **Clone** | **Cat. No.** |
| **Non-immune panel** | | |
| BV421 CD140a (1:100) | APA5 | 135923, BioLegend |
| BUV395 Ep-CAM (1:100) | G8.8 | 740281, BD Biosciences |
| BV605 CD31 (1:200) | 390 | 102427, BioLegend |
| BUV496 CD45 (1:100) | 30-F11 | 749889, BD Biosciences |
| **Immune panel** | | |
| BUV496 CD45 (1:100) | 30-F11 | 749889, BD Biosciences |
| AF488 CD45R/B220 (1:100) | RA3-6B2 | 103225, BioLegend |
| AF532 Ly-6C (1:100) | ER-MP20 | NB100-65413AF532, Novus |
| BUV737 Ly-6G (1:100) | 1A8 | 741813, BD Biosciences |
| BV510 CD11b (1:100) | M1/70 | 101245, BioLegend |
| BV711 CD64 (1:100) | X54-5/7.1 | 139311, BioLegend |
| BV650 CD3 (1:100) | 17A2 | 100229, BioLegend |
| BV570 CD4 (1:100) | RM4-5 | 100541, BioLegend |
| PerCP-eFluor 710 CD8a (1:100) | 53-6.7 | 46-0081-82, Thermo Fisher Scientific |
| Vioblue CD11c (1:100) | N418 | 130-102-797, Miltenyi Biotec |
| BV785 MHCII (1:250) | M5/114.15.2 | 107645, BioLegend |
| AF700 NK-1.1 (1:100) | PK136 | 108729, BioLegend |
| APC-Cy7 F4/80 (1:100) | BM8 | 123117, BioLegend |
| **Intracellular antibody panel** | | |
| PE-Cy7 Bcl-xl (1:200) | 54H6 | 81965, Cell Signaling Technology |
| PE H2AX (1:200, 1:500 for immune panel) | N1-431 | 562377, BD Biosciences |
| Purified recombinant p16 (1:200) (conjugated to AF647) | 2D9A12 | ab54210, Abcam |
| Alexa Fluor™ Antibody Labeling Kits | | A-20186, Thermo Fisher Scientific |

**Supplementary table 5: Extracellular and Intracellular antibody panels used to stain mice intestines**

| Blood | | |
|---|---|---|
| Extracellular antibody panel | | |
| Antibody dilution | Clone | Cat. No. |
| BV605 CD31 (1:200) | 390 | 102427, BioLegend |
| BUV496 CD45 (1:100) | 30-F11 | 749889, BD Biosciences |
| AF488 CD45R/B220 (1:100) | RA3-6B2 | 103225, BioLegend |
| AF532 Ly-6C (1:100) | ER-MP20 | NB100-65413AF532, Novus |
| BUV737 Ly-6G (1:100) | 1A8 | 741813, BD Biosciences |
| BV510 CD11b (1:100) | M1/70 | 101245, BioLegend |
| BV711 CD64 (1:100) | X54-5/7.1 | 139311, BioLegend |
| BV650 CD3 (1:100) | 17A2 | 100229, BioLegend |
| BV570 CD4 (1:100) | RM4-5 | 100541, BioLegend |
| PerCP-eFluor 710 CD8a (1:100) | 53-6.7 | 46-0081-82, Thermo Fisher Scientific |
| Vioblue CD11c (1:100) | N418 | 130-102-797, Miltenyi Biotec |
| BV785 MHCII (1:250) | M5/114.15.2 | 107645, BioLegend |
| AF700 NK-1.1 (1:100) | PK136 | 108729, BioLegend |
| APC-Cy7 F4/80 (1:100) | BM8 | 123117, BioLegend |
| Intracellular antibody panel | | |
| PE-Cy7 Bcl-xl (1:200) | 54H6 | 81965, Cell Signaling Technology |
| PE H2AX (1:100) | N1-431 | 562377, BD Biosciences |
| Purified recombinant p16 (1:200) (conjugated to AF647) | 2D9A12 | ab54210, Abcam |
| Alexa Fluor™ Antibody Labeling Kits | | A-20186, Thermo Fisher Scientific |

**Supplementary table 6: Extracellular and Intracellular antibody panels used to stain mice blood**

| hPBMCs | | |
|---|---|---|
| **Extracellular antibody panel** | | |
| **Antibody dilution** | **Clone** | **Cat. No.** |
| BUV496 CD45 (1:100) | HI30 | 750179, BD Biosciences |
| AF488 CD19 (1:100) | HIB19 | 302219, BioLegend |
| BUV737 CD69 (1:100) | FN50 | 612817, BD Biosciences |
| Vioblue CD3 (1:100) | BW264/56 | 130-113-695, Miltenyi Biotec |
| AF700 CD4 (1:100) | SK3 | 344621, BioLegend |
| BV510 CD8 (1:100) | SK1 | 344731, BioLegend |
| BV421 CD11c (1:100) | Bu15 | 337225, BioLegend |
| BV650 CD16 (1:100) | 3G8 | 302041, BioLegend |
| BV711 CD56 (1:100) | HCD56 | 318335, BioLegend |
| PerCP-eFluor 710 CD31 (1:100) | WM59 | 46-0319-42, Thermo Fisher Scientific |
| BV605 HLA-DR (1:100) | L243 | 307639, BioLegend |
| BUV395 CD14 (1:100) | MφP9 | 563561, BD Biosciences |
| BUV615 Lag-3 (1:100) | T47-530 | 752362, BD Biosciences |
| BV785 CTLA-4 (1:50) | BNI3 | 369623, BioLegend |
| **Intracellular antibody panel** | | |
| PE-Cy7 Bcl-xl (1:1000) | 54H6 | 81965, Cell Signaling Technology |
| PE H2AX (1:1000) | N1-431 | 562377, BD Biosciences |
| AF647 CDKN2A/p16INK4a (1:1000) | EPR1473 | ab192054, Abcam |

**Supplementary table 7: Extracellular and Intracellular antibody panels used to stain hPBMCs**
